# Buoyant mass reveals distinct T cell states predictive of checkpoint response

**DOI:** 10.1101/2024.09.26.615179

**Authors:** Jiaquan Yu, Ye Zhang, Sarah M. Duquette, Robert J. Kimmerling, James C. Kostas, Krystal K. Lum, Rachel LaBella, Selim Olcum, Madeleine Vacha, Steve Wasserman, Reginald Aikins, Katie Katsis, Thomas R. Usherwood, Sonia Cohen, Teresa Dinter, Aleigha Lawless, Tatyana Sharova, Mark Stevens, Gianfranco L. Yee, Weida Wu, Stefani Spranger, Teemu P. Miettinen, Ileana M. Cristea, Genevieve M. Boland, Scott R. Manalis

**Affiliations:** Koch Institute for Integrative Cancer Research, Massachusetts Institute of Technology, Cambridge, MA 02139, USA; Department of Biological Engineering, Massachusetts Institute of Technology, Cambridge, MA 02139, USA; Travera, Medford, MA, USA; Department of Molecular Biology, Princeton University, Lewis Thomas Laboratory, Princeton, NJ 08544, USA; Department of Biology, Massachusetts Institute of Technology, Cambridge, MA, USA; Harvard-MIT Department of Health Sciences and Technology, Institute for Medical Engineering and Science, Massachusetts Institute of Technology, Boston, MA, USA; Massachusetts General Hospital, Krantz Family Center for Cancer Research, Boston, MA, USA; Harvard Medical School, Boston, MA, USA

## Abstract

T cells are central to immune defense, yet existing molecular and phenotypic assays do not fully capture a cell’s intrinsic immune potential. Here we show that a single physical property, buoyant mass, reveals hidden heterogeneity within phenotypically similar, resting CD8^+^ T cells. Using suspended microchannel resonator measurements, we identify two distinct populations: “light” cells, enriched for mitochondrial content but prone to delayed activation and exhaustion, and “heavy” cells, biosynthetically poised for proliferation and memory formation. In patients with melanoma receiving immune checkpoint blockade, pre-treatment buoyant mass profiling of circulating T cells predicted therapeutic response with an accuracy comparable with standard tumor-derived biomarkers. Our findings establish buoyant mass as a label-free, stimulation-independent measure of systemic T cell fitness, providing a rapid and broadly applicable framework for immune profiling and response prediction in cancer and beyond.

## Introduction

T cells are central to immune surveillance, with their activation, proliferation, and differentiation into distinct functional states governed by a complex integration of stimulatory and inhibitory cues. Existing assays that assess molecular markers, gene expression, or cytokine secretion capture specific aspects of T cell biology but may not fully reflect its integrated functional potential. This limitation is particularly relevant for cancer immune checkpoint blockade therapy. Currently used biomarkers^1,2,3^, such as tumor mutational burden, PD-L1 expression, and microenvironmental features, have informed patient stratification, but they primarily focus on correlative markers in the tumor microenvironment rather than systemic immune competence. Even when peripheral blood is examined^4,5^, current approaches emphasize composition (cell counts, phenotypes, or protein levels) rather than intrinsic functional potential. As a result, there is no widely adopted method to directly assess the functional capacity of circulating T cells, despite their central role in checkpoint inhibitor efficacy.

Biophysical properties of cells, such as size, density, and mass, offer an integrative view of cellular state. Buoyant mass, in particular, reflects the combined effects of cell volume and composition, including protein and organelle content^6,7^. Suspended microchannel resonator (SMR) technology enables buoyant mass to be measured at the single-cell level with picogram precision, and prior study has shown its utility in distinguishing proliferating from quiescent cells^8,9^. Whether buoyant mass reveals clinically relevant variation in unstimulated, phenotypically similar T cells remains unknown.

Here we report that buoyant mass uncovers a previously hidden layer of heterogeneity within resting CD8^+^ T cells, dividing them into two populations with distinct mitochondrial programs, proliferative kinetics, and exhaustion propensity. These biophysical states highlight the link between buoyant mass and T cell function, supporting its use to stratify immune competence in ways that directly predict clinical outcomes. In a cohort of patients with melanoma receiving immune checkpoint blockade, pre-treatment buoyant mass profiling of circulating T cells predicted therapeutic response with an accuracy comparable with tumor mutational burden, a standard tumor-derived biomarker. Our findings establish buoyant mass as a simple, label-free measure of systemic T cell fitness, offering a broadly applicable framework for immune monitoring and therapeutic response prediction.

## Results

We applied the SMR to profile both resting (i.e., unactivated), and for comparison, activated CD8^+^ T cells (**Fig. 1a**). Among resting T cells, we observed a bimodal distribution of buoyant mass: a “light” population (5–7.5 pg) and a “heavy” population (>7.5 pg) with a clear upper threshold of 16 pg (**Fig. 1b, c**). This distribution is distinct from that of activated T cells, which can reach buoyant masses of up to 60 pg (**Fig. 1b**)^8^. Upon in vitro activation of mouse CD8^+^ T cells with anti-CD3/CD28 stimulation^10,11^, we analyzed the cells after 24 hours using a fluorescence-coupled SMR and the proliferation marker Ki67. All Ki67^+^ (dividing) cells fell above 12 pg, largely exceeding the heavy population within unactivated cells (**Fig. 1b**). In contrast, no Ki67^+^ cells were detectable prior to stimulation. We confirmed the bimodal buoyant mass distribution of resting CD8^+^ T cells across 38 independent mouse spleen samples. Kernel density analysis identified a consistent antimode (the valley between peaks) at ∼7.5 pg (**Fig. 1c**). Each distribution could be quantitatively characterized by three features: the percentage of light cells (**Fig. 1d**), the mean mass of light cells (**Fig. 1e**), and the mean mass of heavy cells (**Fig. 1f**). This unexpected bimodal mass distribution in resting CD8^+^ T cells suggests that biophysical heterogeneity may underlie unappreciated layers of T cell physiology.

**Fig. 1.**
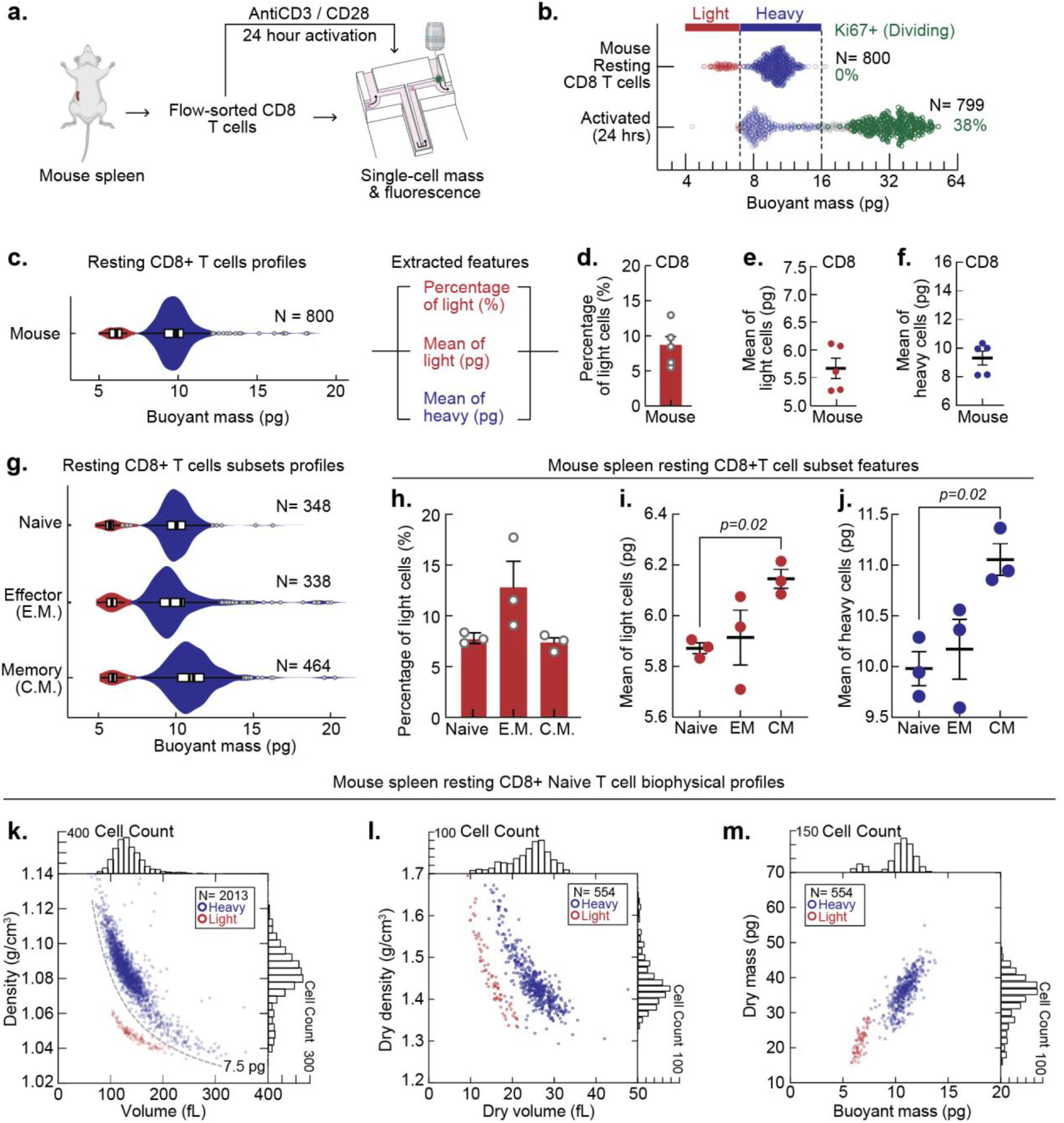
Mouse CD8+ T cell samples consistently exhibit a bimodal buoyant mass distribution. **a**, Schematic of the workflow for sorting CD8+ T cells from mouse spleen samples by flow cytometry for resting or activated measurements using the fluorescent coupled SMR for single-cell mass profiling and fluorescent pairing. **b**, Buoyant mass profiles of CD8+ T cells before activation and after 24 hrs of activation. Cells actively expressing Ki67 at the time of measurement are indicated in green. **c**, Mass distribution of mouse CD8+ T cell samples displaying light (5-7.5 pg) in red and heavy (>7.5 pg) in blue. Bimodal distributions are annotated with median +/-50% for each mode with a dotted line indicating the antimode. **d**, The percentage of light (red) T cells within a CD8+ T cell mouse sample. Mean ± s.e.m. are shown. **e-f**, Mass mean of **e**, light (red) and **f**, heavy (blue) populations within CD8 T cells. **g**, Buoyant mass profiles of naive (CD44-,CD62L+), effector memory (EM, CD44+,CD62L-), and central memory (CM, CD44+,CD62L+) T cells. Color indicates mass subpopulation with light in red and heavy in blue. Bimodal distributions are annotated with median +/- 50% for each mode with a dotted line indicating the antimode. **h**, The percentage of light (red) and heavy (blue) T cells within the three CD8+ T cell subpopulations (n=3). Mean ± s.e.m. are shown. **i-j**, Mass mean of **i**, light (red) and **j**, heavy (blue) populations within CD8 T cell subsets. Statistical analysis was performed using one-way ANOVA with mean value multi-comparison. **k**, Scatter plot of density (g/cm^3^) vs volume (fL) (dashed line indicates iso-buoyant mass), **l**, dry density (g/cm^3^) vs dry volume (fL). (dashed line indicates iso-dry mass), and **m**, dry mass (pg) vs buoyant mass (pg) for naïve CD8+ T cells from mouse spleen.

To explore whether these light and heavy cell populations correspond to distinct T cell differentiation states, we sorted live mouse CD8^+^ T cells into three canonical subsets: naïve (CD44^−^CD62L^+^), effector memory (CD44^+^CD62L^−^), and central memory (CD44^+^CD62L^+^) (**Extended Data Fig. 1a**)^12,13^. Flow sorting purity exceeded 99.8%, as validated by single-cell RNA sequencing (**Extended Data Fig. 2b,d**). Mass features were extracted from buoyant mass profiles of each subset revealing the continued presence of both light and heavy populations within all three groups, as well as a difference in the median of the heavy population between subsets (**Fig. 1g**). Intriguingly, the mean buoyant mass of heavy population alone was sufficient to distinguish central memory cells from both naïve and effector memory subsets (**Fig. 1j**), while percent light and median buoyant mass of the light population could not (**Fig. 1h,i**). This highlights the link between T cell buoyant mass and cell function suggesting buoyant mass features as potential biomarkers of T cell functional capacity.

**Fig. 2.**
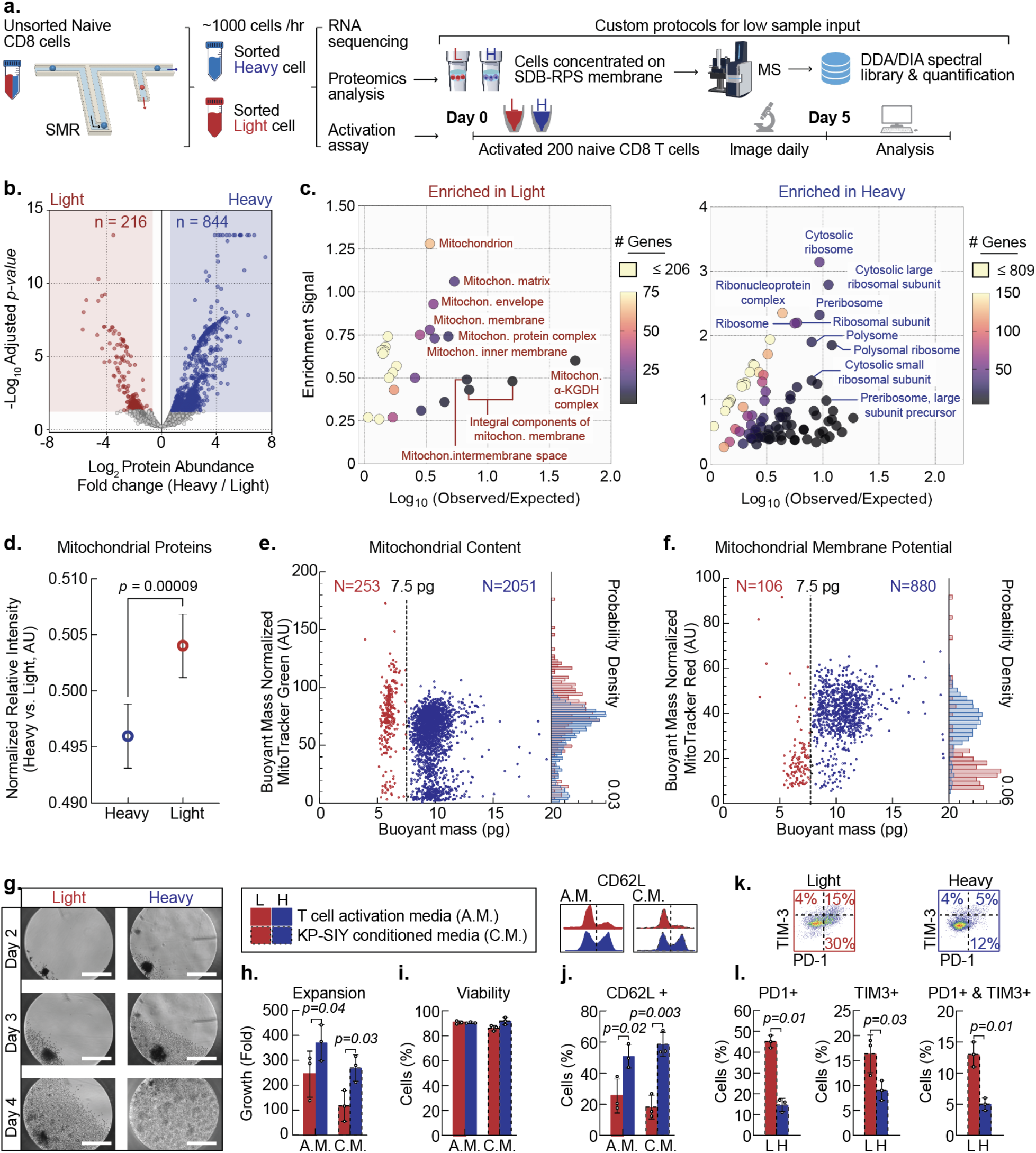
Light and heavy T cells differ in mitochondrial content. **a**, Schematic of mass-based single-cell sorting platform via SMR for subsequent transcriptional and functional analysis. The sorting accuracy is over 98%. Schematic of the workflow for proteomic assessment of heavy and light T cells by mass spectrometry. **b**, Scatter plot of relative protein abundances (heavy over light cells). Differentially abundant proteins are colored (adjusted p-value ≤ 0.05 and log2 fold change ≤ -0.58 (red, light cells) or ≥ 0.58 (blue, heavy cells). **c**, Gene ontology cellular component analysis of differentially abundant proteins in **b** using STRING^34^. **d**, Comparison of all identified proteins with reported mitochondrial localization in heavy versus light cells (unpaired t test with Welch’s correction, *p*-value < 0.0001). Solid lines indicate upper and lower quartiles. Dotted lines indicate the median. **e**, Mass normalized MitoTracker Green fluorescent signal vs buoyant mass (pg) measured for CD8+ T cells. **f**, Mass normalized MitoTracker deep red FM fluorescent signal vs buoyant mass (pg) measured for CD8+ T cells. **g**, Brightfield images of T cells during activation with KP-SIY conditioned media (C.M.) on days 2, 3, and 4. The scale bar is 300 μm. **h-i**, Bar plot of total cell cultured in T cell activation media (A.M.) or C.M. **h**, expansion and **i**, viability calculated by the fold change of cell counts on day 6 to the number of cells seeded plotted for light and heavy. Data are shown as mean ± s.e.m., representative of three independent experiments. Statistical analysis was performed using two-tailed paired t-tests with Welch’s correction. **j-l**, Analysis via Amnis imagestream to analyze **j**, CD62L, **l**, PD-1, and TIM3 expression on day 6 of activation. **k**, Representative flow plots of PD-1 and TIM3 gating strategy for activated heavy and light. **h-j**, & **l**, Data are shown as mean ± s.e.m., representative of three independent experiments. Statistical analysis was performed using two-tailed unpaired t-tests with Welch’s correction.

### Light and heavy T cells are distinguished only by buoyant mass

The bimodal distribution of resting CD8^+^ T cells was robust across experiments, prompting us to test whether standard characterization methods could detect this heterogeneity. Flow cytometry analysis using forward and side scatter showed a single, compact cluster with no evidence of two populations (**Extended Data Fig. 3a**). Sorting cells from the high- and low-scatter quadrants and re-measuring buoyant mass revealed that both fractions contained overlapping light and heavy cells (**Extended Data Fig. 3b)**, confirming that scatter-based size and granularity measures fail to capture the bimodal phenotype. Imaging flow cytometry likewise showed no separation in basic morphological parameters such as cell area, perimeter, or aspect ratio (**Extended Data Fig. 3c**). Thus, despite clear functional differences, resting CD8^+^ T cells appear homogeneous by conventional optical assays, with bimodality apparent only through buoyant mass profiling.

**Fig. 3.**
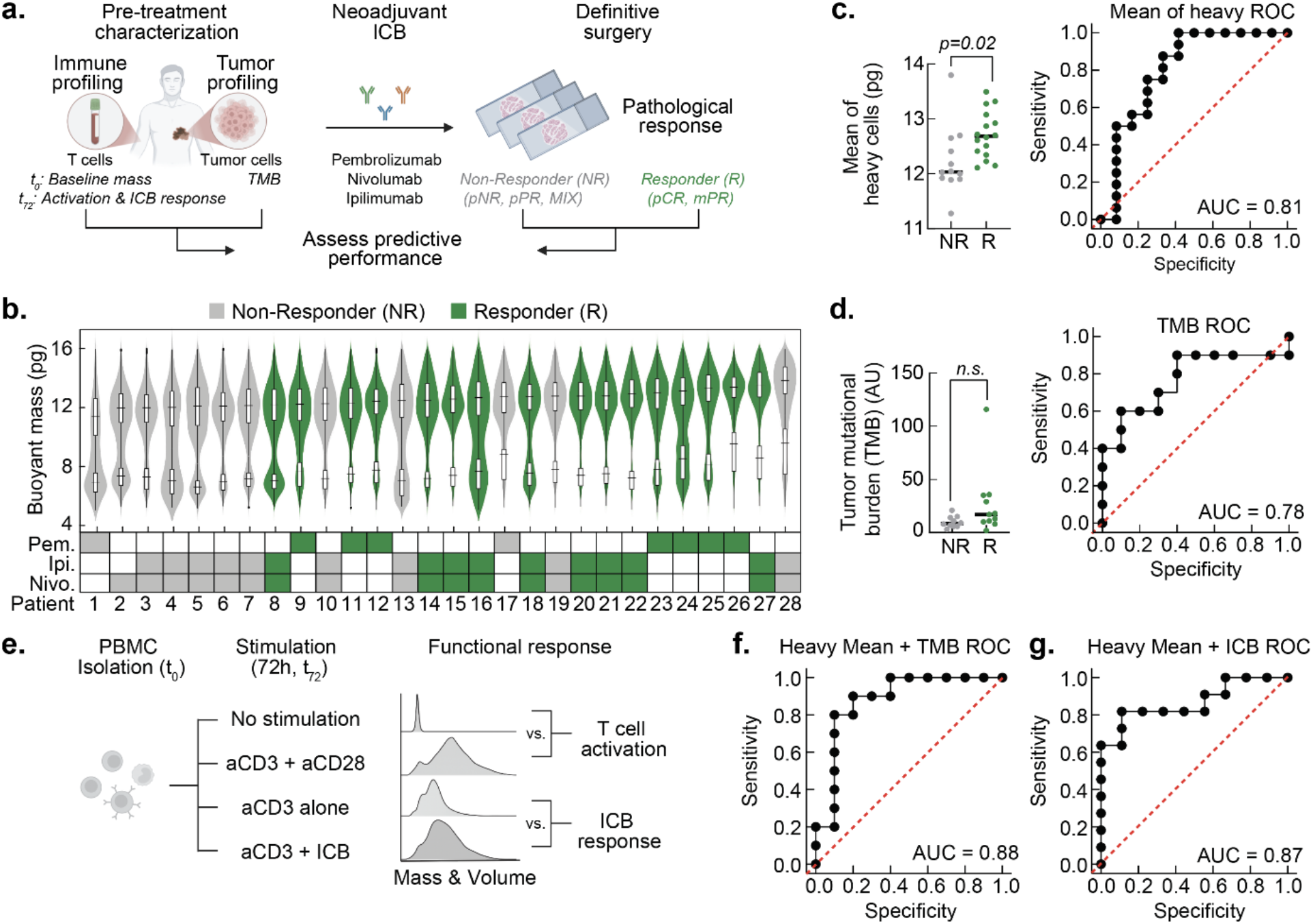
Biophysical profiling of peripheral T cells predicts response to ICB in melanoma patients. **a**, Schematic of neoadjuvant ICB treatment in melanoma study. **b**, Buoyant mass distributions of pretreatment peripheral blood T cells from 28 patients, ordered by heavy mean and colored by response to ICB therapy. **c**, Comparison of mean heavy T cell mass between responders (n=16) and non-responders (n=12). Responders exhibited significantly greater heavy-cell mass (Mann-Whitney U test, p < 0.01), yielding an AUC of 0.81. **d**, Comparison of tumor mutational burden (TMB) between responders and non-responders (n=21 patients). TMB is quantified as the number of mutations per megabase. Differences were not statistically significant (Mann-Whitney U test, p > 0.05). **e**, Ex vivo T cell activation with anti-CD3 alone, anti-CD3/CD28, or anti-CD3+ICB. ICB response was quantified as the incremental mass gain between anti-CD3 and anti-CD3+ICB conditions. **f,g**, Assessment of heavy mean combined with TMB and with ICB response respectively to improve predictive power.

Since buoyant mass depends on cell volume and buoyant density, which in turn depend on dry volume and dry density, respectively^14,15^, we utilized an established method to measure these additional biophysical parameters (**Extended Data Note 1**). Remarkably, naive CD8+ T cells isolated from mouse spleen show an overlap of light and heavy for both volume and buoyant density, indicating that neither volume nor buoyant density alone accounts for the bimodality observed in T cell buoyant mass (**Fig. 1k**). Yet, when plotting buoyant density versus volume for each individual cell, the two distinct populations are readily apparent, as anticipated by the iso-buoyant mass contour (**Fig. 1k**). Similarly, the light and heavy populations also overlap in terms of dry volume and dry density (**Fig. 1l**), but when plotted versus each other, the two populations are distinct. In contrast, the iso-dry mass contour reveals overlap between these two populations. This is more readily apparent when dry mass is plotted versus buoyant mass, showing clear bimodality for the latter but not the former (**Fig. 1m**). Cell dry mass has long been used as a metric for cell growth because it integrates all cellular components, including protein, nucleic acid, and lipid content. Although, in prior growth studies buoyant and dry mass generally lead to similar qualitative conclusions^14^, they reflect fundamentally different properties of the cell. For cells in media where the density is close to that of water, the relationship between the cell’s buoyant and dry mass is governed by its dry density (**Extended Data Note 1**). Taken together, these results demonstrate that the bimodal buoyant mass profile of resting CD8^+^ T cells can only be resolved through buoyant mass measurements.

### Light and heavy naive T cells differ in mitochondrial content, expansion, and exhaustion potential

To investigate functional differences between light and heavy naive T cells, we developed an enhanced SMR chip that enables the collection of distinct mass populations using a predefined buoyant mass cutoff **(Fig. 2a)**. Specifically, we integrated a binary branch into the silicone-etched SMR flow channel, allowing on-chip sorting of up to 1,000 cells per hour. Our platform utilizes three independent pressure regulators to control the single-cell loading channel and two output sorting channels **(Extended Data Fig. 4a)**. This system represents, to our knowledge, the first demonstration of sorting live cells by buoyant mass in a flow-cytometry-like, bin-gated manner.

To probe the molecular basis of light and heavy naïve T cell populations, we performed bulk RNA sequencing (RNA-seq) on cells from three mice sorted by buoyant mass. RNA-seq identified 102 differentially expressed genes (DEGs) between light and heavy cells (adjusted p < 0.05; log_2_ fold change > 1.5) (**Extended Data Fig. 5a**). Interestingly, highly expressed DEGs (>104) were predominantly mitochondrial, with six of the top eight DEGs originating from the mitochondrial genome (**Extended Data Fig. 5b**). Quantification of mitochondrial RNA fractions revealed that light cells contained 3.9% more mitochondrial RNA than heavy cells (p = 0.049) (**Extended Data Fig. 5c**). Additionally, hierarchical clustering of mitochondrial genes confirmed their upregulation in light cells, effectively distinguishing the light and heavy populations (**Extended Data Fig. 5d**). While elevated mitochondrial RNA levels can sometimes reflect cell stress or damage, gene set enrichment analysis revealed no significant differences in apoptosis, autophagy, mitophagy, or cell cycle pathways (**Extended Data Fig. 6**). These results suggest that the observed mitochondrial differences reflect a distinct transcriptional program, rather than generalized stress responses.

To complement our transcriptomic analysis, we performed mass spectrometry–based proteomics on the light and heavy naïve T cells. Unlike RNA-seq, proteomics directly quantifies protein abundance and post-translational modifications, providing a more immediate snapshot of cellular state^16,17^. Conducting proteomics on buoyant-mass sorted T cells posed challenges due to low cell numbers (∼1,000 per mouse) and minimal protein content. To overcome this, we developed a low-input sample preparation workflow that enabled complete processing (from lysis to peptide cleanup) in minimal volumes (**Fig. 2a, Extended Data Fig. 7a**). Samples were processed using micropipette tips packed with a styrene-divinylbenzene membrane, which allowed effective detergent removal while preserving protein recovery for LC–MS/MS analysis. To maximize protein detection, we constructed a hybrid mass spectral library using both data-dependent (DDA) and data-independent acquisition (DIA) modes. DIA-PASEF data from light and heavy samples were then analyzed using this custom library. Proteomic profiles clustered primarily by mouse (**Extended Data Fig. 7b**), mirroring our RNA-seq results and underscoring the biological heterogeneity across individuals.

With this optimized workflow, we identified ∼5,000 unique proteins across all samples. Differential abundance analysis (padj ≤ 0.05; log_2_FC ≥ 0.58) revealed 216 proteins enriched in light cells and 844 in heavy cells (**Fig. 2b, Supplementary Table 1**). Consistent with RNA-seq results, light cells showed strong enrichment for mitochondrial proteins, including components of the matrix, membrane, and α-ketoglutarate dehydrogenase complex (**Fig. 2c**). In contrast, heavy cells were enriched for cytosolic ribosomal proteins, suggesting enhanced translational capacity. These findings point to a bioenergetic distinction where light T cells may experience mitochondrial overactivation and metabolic stress, while heavy T cells appear more biosynthetically primed. Indeed, annotation-based quantification showed significantly higher mitochondrial protein content in light cells (p < 0.0001) (**Fig. 2d**), mirroring their increased mitochondrial RNA levels. To functionally validate these differences, we stained naïve CD8^+^ T cells with MitoTracker Green (total mitochondrial mass) and MitoTracker Deep Red FM (membrane potential). Light cells exhibited higher mitochondrial mass but lower membrane potential, suggesting increased mitochondrial biogenesis coupled with impaired function (**Fig. 2e, f**). These results are consistent with our transcriptomic, proteomic, and functional modalities.

Given their reduced mitochondrial membrane potential, we hypothesized that light T cells would exhibit impaired proliferative and differentiation capacity. To test this, we activated buoyant mass–sorted cells using two protocols simulating conventional T cell activation and under tumor conditioned media expansion (**Fig. 2a**). Although light and heavy T cells formed clusters of similar size initially, heavy cells expanded more rapidly and outnumbered light cells by day 6, despite comparable viability (**Fig. 2g–i**). Light cells nonetheless accumulated buoyant mass (**Extended Data Fig. 8c**), suggesting delayed but intact proliferative response. We next evaluated CD62L re-expression, a marker of memory potential. A significantly smaller fraction of light T cells reacquired CD62L compared to heavy cells, a difference observed across both activation conditions (**Fig. 2j**). These findings suggest that light T cells may be less poised to differentiate into central memory T cells. To investigate whether this reflects early exhaustion, we examined expression of PD-1 and TIM-3. Light cells showed twice the frequency of PD-1^+^, TIM-3^+^, or double-positive cells compared to heavy cells (**Fig. 2k-l**). Together, these data indicate that light T cells follow a distinct functional trajectory with delayed activation, limited memory formation, and early exhaustion.

### T cell biophysical profiling predicts response to immune checkpoint blockade

The distinct proliferative, metabolic, and differentiation trajectories observed between light and heavy T cells suggest that buoyant mass may serve as an integrated indicator of systemic immune competence, capturing dimensions of T cell fitness not readily assessed by molecular markers alone. We therefore evaluated whether these biophysical signatures could predict clinical response to immune checkpoint blockade (ICB) therapy from T cells within peripheral blood. We studied a cohort of 28 patients with locally advanced melanoma receiving standard-of-care neoadjuvant ICB, including combination ipilimumab and nivolumab (n = 18), pembrolizumab monotherapy (n = 9), or nivolumab monotherapy (n = 1) (**Fig. 3a**). Peripheral blood was collected prior to treatment to measure baseline T cell buoyant mass and to perform ex vivo functional profiling. We compared these immune profiles with pathological response outcomes, assessed at the time of definitive surgery. Pathological response has been shown to correlate strongly with long-term outcomes in melanoma^18^, and patients in this study did not receive concomitant cytotoxic chemotherapy or radiation, enabling a direct evaluation of ICB efficacy. In this cohort, 16 patients responded to treatment, including 14 with a pathological complete response (pCR) and 2 with a major pathological response (MPR). The remaining 12 patients were non-responders, including 9 with pathological non-response (pNR), 2 with partial response (pPR), and 1 with a mixed response across disease sites. This response rate was consistent with recent neoadjuvant ICB trials in resectable stage III melanoma^19^ and provided a clinically meaningful framework for assessing biomarker performance.

In pre-treatment blood from 28 patients, we measured >7,000 cells per patient on average (**Extended Data Fig. 10**) and consistently observed a bimodal T cell distribution (**Fig. 3b**). Building on our earlier observation that distinct T cell subsets exhibit characteristic heavy-cell mass profiles (**Fig. 1j**), we next asked whether variation in heavy T cell mass correlated with clinical outcome. Responders to ICB had a higher mean heavy-T-cell mass (Δ=0.48 pg) than non-responders (p = 0.02; **Fig. 3c**), with per-sample standard error of the mean (SEM) ranging from 0.012 to 0.069 pg (**Extended Data Fig. 10**). This single metric alone distinguished the two groups with high accuracy (AUC = 0.81; **Fig. 3c**). In contrast, when the full T cell mass distribution, including both light and heavy populations, was analyzed, predictive performance declined markedly (AUC = 0.71), and the difference in mean T cell mass was no longer significant (p > 0.05; **Extended Data Fig. 11a**). These results highlight the importance of focusing specifically on subtle buoyant mass shifts within the heavy population, a distinction made possible by the precision of the SMR. Moreover, responders exhibited a lower standard deviation in heavy T cell mass (**Extended Data Fig. 11b**), suggesting a more uniform biophysical phenotype in this group. Collectively, these findings indicate that the heavy T-cell buoyant-mass profile serves as a biomarker of immune state and predicts therapeutic response.

To benchmark biophysical profiling, we compared its performance with tumor mutation burden (TMB), a standard clinical biomarker for ICB response prediction. Among 20 patients with available TMB data, TMB demonstrated an AUC of 0.78 when distinguishing responders from non-responders (**Fig. 3d**), though the difference in TMB values between responders and non-responders did not reach statistical significance (p>0.05, **Fig. 3d**). This performance is consistent with previous work demonstrating mixed utility of TMB as a biomarker for ICB response^20^.

In addition to heavy T cell mass measurements, we assessed whether functional biophysical readouts that capture T cell responsiveness *ex vivo* could further discriminate responders from non-responders. For a subset of patients, we quantified two activation-dependent metrics: (1) the change in buoyant mass following stimulation with anti-CD3/CD28, as a proxy for general T cell activation, and (2) the incremental shift in single-cell mass and volume induced by addition of ICB drug during activation, reflecting checkpoint-specific responsiveness (**Fig. 3e**). Although T cell activation alone demonstrated modest discriminatory power (AUC = 0.64; **Extended Data Fig. 11c**), the checkpoint-specific activation more robustly distinguished responders from non-responders (AUC = 0.78; **Extended Data Fig. 11d**). The biophysical response to ICB was significantly different in responders versus non-responders (p = 0.036; **Extended Data Fig. 11d**), suggesting that checkpoint sensitivity is reflected in T-cell biophysics.

Building on recent evidence that multimodal data integration improves ICB response prediction^4,21,22^, we next asked whether combining these biomarkers could further enhance predictive accuracy. Indeed, integrating heavy T cell mass with TMB (n = 20) improved response classification performance (AUC = 0.88; **Fig. 3f**), as did the combination of heavy T cell mass with checkpoint-specific activation response (AUC = 0.87; **Fig. 3g**). Both multimodal approaches outperformed any single marker alone. These findings show the complementary nature of tumor-centric and immune-intrinsic biomarkers and support the value of incorporating T cell mass profiling into multiparametric strategies for predicting response to ICB.

## Discussion

The search for reliable predictors of immunotherapy response has increasingly turned toward computational integration of diverse molecular and clinical features. Recent efforts such as SCORPIO, a machine learning framework trained on thousands of patients using routine clinical labs and metadata, have demonstrated the power of large-scale, data-driven models to forecast checkpoint inhibitor outcomes across cancer types^4^. Similarly, LORIS, a logistic regression–based score derived from a handful of clinical, genomic, and pathologic variables, provides interpretable probability estimates of response and survival, outperforming TMB and PD-L1 expression in multiple cohorts^21^. These approaches demonstrate the value of statistical learning in integrating complex variables to stratify patients. By contrast, our method does not rely on computational feature aggregation or tumor-intrinsic metrics. Instead, we leverage a single physical property (buoyant mass of unstimulated circulating T cells) to capture integrated aspects of immune competence. Remarkably, in melanoma, a tumor type known to be highly immunogenic, this systemic immune measure alone achieves predictive accuracy comparable to or greater than multi-feature ML models, underscoring its potential as a powerful standalone biomarker. At the same time, because peripheral blood profiling is minimally invasive and broadly scalable, these measurements can also be incorporated into multiparametric frameworks in other malignancies, where combining systemic immune metrics with tumor-intrinsic or clinical features may further enhance predictive performance.

Interestingly, our analyses revealed that the mean buoyant mass of heavy T cells in both responders and non-responders was comparable to that of healthy donors, with no significant differences observed (**Extended Data Fig. 11e**). This finding suggests that variation in heavy T cell mass is not simply a reflection of disease-induced alterations but rather may represent innate inter-individual differences in immune constitution. Such baseline variability highlights the need for longitudinal and cross-disease studies to disentangle trait-like versus state-like determinants of T cell buoyant mass, and to clarify whether individuals with inherently higher or more stable heavy T cell profiles are predisposed to mount superior responses to immunotherapy.

Finally, while immune checkpoint blockade provides a compelling clinical application, the implications of biophysical immune profiling may extend beyond oncology. In autoimmune disease, organ transplantation, chronic infection, and aging, there is a need for quantitative metrics of immune competence that are both functional and practical. Most existing assays rely on stimulated responses or molecular surrogates that can be costly, variable, and time-consuming to interpret. In contrast, the buoyant mass assay presented here requires only peripheral blood T cells, no stimulation or labeling, and can be completed in under 20 minutes on a platform already validated in a CLIA-certified setting. This operational simplicity, combined with its ability to stratify functional immune states without prior knowledge of disease context, positions buoyant mass as a broadly applicable, real-time biomarker of systemic immunity. The discovery that a single physical property can reveal hidden dimensions of T cell heterogeneity suggests that biophysical profiling may serve as a foundational layer for assessing immune fitness across a wide range of diseases and therapeutic interventions.

## Methods

### Mice

All mice used in Figs. 1-5 were bred and maintained under specific pathogen-free conditions at the Koch Institute Animal facility. Animal procedures were approved by the Massachusetts Institute of Technology Committee on Animal Care (CAC), Division of Comparative Medicine (DCM). Animals were housed on hardwood chip bedding, with a 12/12 hour light-dark cycle at a temperature of 70°F +/− 2 and humidity of 30–70%. These mice were maintained on a mixed C57BL/6; 129/Sv background. Mice were 12-20 weeks old at the time of experimentation, unless otherwise described.

### Mouse Spleen processing

Mice were euthanized via CO_2_ asphyxiation. The spleen was removed and mashed through a 70 μm filter to create a single cell suspension. The spleen sample was then enriched for CD8+ T cells via the EasySep Mouse CD8+ T cell Isolation Kit (StemCell, 19853) following the manufacturer’s protocol. The CD8+ T cell enriched sample was then stained with FC Block (101302), eBioscience Fixable Viability Dye (APC-Cy7, 65-0865-14), CD19 (APC-Cy7, 47-0193-80), CD4 (APC-Cy7, 47-0042-80), NK1.1 (APC-Cy7, 47-5941-80), CD8 (BV421, 100725), CD62L (APC, 17-0621-81), and CD44 (FITC, 11-0441-81) for fluorescent activated cell sorting of CD8+ T cells and T cell subpopulations.

### Human PBMC processing

Apheresis leukoreduction collars from anonymous healthy platelet donors were obtained from the Brigham and Women’s Hospital Specimen Bank under an Institutional Review Board–exempt protocol. Human peripheral blood mononuclear cells (PBMCs) were isolated via density gradient centrifugation (Lymphoprep, StemCell Technologies Inc, 07801). PBMC samples were then stained with FC Block (422301), eBioscience Fixable Viability Dye (APC-Cy7, 65-0865-14), CD19 (APC-Cy7, 363009), CD4 (APC-Cy7, 47-0049-41), CD56 (APC-Cy7, 47-0567-41), CD14 (APC-Cy7, 47-0149-41), CD8 (APC, MHCD0821), CD197(CCR7) (PE, 353203), and CD45RA (PE-Cy7, 25-0458-42) for fluorescent activated cell sorting of CD8+ T cells.

### T cell buoyant mass measurement using SMR

Single-cell buoyant mass was measured using the SMR as previously described^23^. The SMR is a cantilever-based mass sensor with an embedded microfluidic channel. As a cell flows through the channel along the cantilever, the change in resonance frequency detected corresponds to the buoyant mass of the cell. Full measurement details can be found in our previous publication^23^. Prior to a set of measurements, the SMR was cleaned with 0.05% Trypsin-EDTA (Invitrogen, 25300054) for 20 min, followed by 5% bleach for 3 min and then a 5-min rinse with DI-H_2_O, to remove persistent biological debris. After cleaning, the SMR was passivated with 1 mg/mL PLL-g-PEG (Nanosoft Polymers, SKU#11354) in H_2_O for 10 min at room temperature, followed by a 5-min rinse with Flow Cytometry (FACS) Staining Buffer (Rockland, MB-086-0500). All measurements were carried out in FACS buffer at room temperature for no more than 30 minutes. The SMR was briefly washed with the FACS buffer between each sample.

During the measurement, all the samples were loaded into the SMR through 0.005-inch-inner-diameter fluorinated ethylene propylene (FEP) tubing (Idex, 1576L). The fluid flow across the SMR was driven by three independent electronic pressure regulators (MPV1, Proportion Air) and three solenoid valves (S070, SMC). A consistent differential pressure was applied across the SMR to maintain constant shear and data rate for cell measurement. All the regulators, valves and data acquisition were controlled by custom software coded in LabVIEW 2020 (National Instruments).

### T cell density and volume measurement

Cell density and volume were measured using fluorescent exclusion techniques as previously described. To couple single-cell mass and volume measurements, a fluorescent microscope was positioned at the entry to the SMR cantilever. The fluorescence level emitted from the detection region was continuously monitored by a photomultiplier tube (PMT, Hamamatsu, H10722-20). Cells were suspended in PBS +2%FBS with 5 mg/mL FITC-conjugated dextran (Sigma, FD2000S-250MG). When no cell was present at the detection region, the PMT detected a high fluorescence baseline from the fluorescence buffer. As a cell passed through, the fluorescence signal decreased proportionally to the volume of the cell. Immediately after the volume measurement, each single cell flowed through the SMR cantilever and the corresponding buoyant mass was measured.

Buoyant mass is given by:

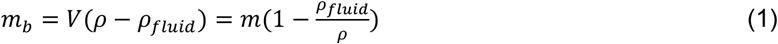

where *V* is the volume of the cell, *m* is the total mass and *ρ* is the density of the cell immersed in a fluid of density *ρ*_*fluid*_. The fluid density of PBS +2%FBS with 5 mg/mL FITC-conjugated dextran is 1.005 g/cm^-3^. Cell density *ρ* is then computed using equation (1).

### T cell dry mass measurement

Cell dry mass and dry density were calculated from SMR measurements as previously described^14^. When measuring cells in PBS, the relationship between buoyant mass and dry mass is closely approximated by” The relationship between buoyant mass and dry mass is given by

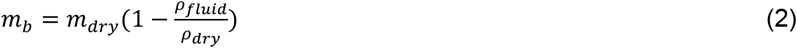

where *ρ*_*fluid*_ is the density of the fluid that the cell is immersed in. To measure dry mass and dry density, the two bypass channels on either side of the SMR cantilever were filled with different fluids, one with an H_2_O-based PBS solution, and the other with a D_2_O-based PBS solution (Sigma, 151882-1KG). Two consecutive buoyant mass measurements were taken of individual T cells; the first was taken with the cell immersed in the H_2_O-based solution, and after a brief period of stopped flow for the intracellular H_2_O to exchange to D_2_O, the second was taken with the cell immersed in a D_2_O-based PBS solution. Given the density difference between H_2_O and D_2_O and the intracellular fluid exchange between the two buoyant mass measurements, equation (1) in these two cases yields

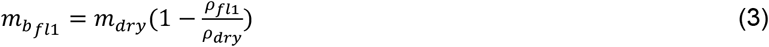

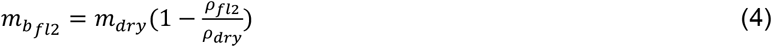

where 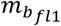 and 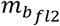 are measured buoyant mass of the cell immersed in H_2_O-based PBS solution (*ρ*_*fl*1_=1.002 g/cm^-3^) and D2O-based PBS (*ρ* _*fl*2_=1.104 g/cm^-3^) respectively. Dry mass and dry density of the cell can be calculated by

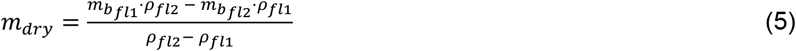

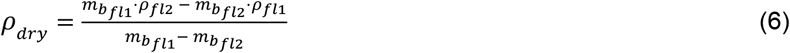

### Mitochondrial staining

Live CD8+ T cells sorted via flow cytometry were stained in PBS with MitoTracker Deep Red FM (Thermofisher, M22426) or MitoTracker Green FM (Thermofisher, M7514) following the manufacturer’s protocol. Cells were incubated with 500 nM MitoTracker for 15 minutes at room temperature. After staining, cells were washed once and immediately analyzed by the fluorescent coupled SMR.

### Single cell RNA sequencing

At least 100k Naive CD8+ T cells sorted on flow cytometry (pre-SMR) and ∼20k Naive CD8+ T cells after flowing through the SMR (post-SMR) were processed for 10x Genomics library preparation using 10x Genomics Chromium Single Cell 3’ RNA assay (v4/GEM-X) following the manufacturer’s protocol using 11 cycles of amplification to generate cDNA. The resulting libraries were quantified and fragment size distribution was assessed using an Agilent Fragment Analyzer and qPCR. Libraries were pooled at equimolar concentrations and sequenced on an Illumina NextSeq 500 platform using 75-cycle High Output Kit (paired-end: R1: 26 nt, i7 index: 8 nt, R2: 57 nt) to achieve an average depth of ∼ 20,000 reads per cell. Cells were processed using CellRanger with default configurations.

### Single cell sequencing data analysis

#### Comparing pre and post SMR

Cells with less than 200 features detected and genes detected in fewer than 3 cells were filtered out. Data was normalized using the NormalizeData function in Seurat. Two-dimensional embeddings were generated using uniform manifold approximation and projection (UMAP). Clusters were identified using the FindClusters function in Seurat.

#### Purity of preSMR

Cells with less than 300 and greater than 5000 features detected or with greater than 20% mitochondrial gene counts and genes detected in fewer than 3 cells were filtered out. Data was normalized using the NormalizeData function in Seurat. Two-dimensional embeddings were generated using uniform manifold approximation and projection (UMAP). Clusters were identified using the FindClusters function in Seurat. DEG analysis was performed for each cluster using the FindMarkers function through presto. Number of contaminating NK cells, neutrophils, B cells, and monocytes was determined by calculating the number of cells expressing 3 or more markers of each indicated cell type. List of identifying markers used is available in **Supplementary Table 4**.

### Mass-based cell sorting

T cells were sorted based on buoyant mass measured by the SMR and observed by brightfield microscopy as they were controlled by standard pressure-driven fluidic components (**Extended Data Fig. 4a**). Prior to sorting, cell concentrations were quantified utilizing a Coulter Counter (Beckman Inc.) and subsequently adjusted to a density of 100,000 cells/mL in PBS if measured concentration exceeded this value. The sorting parameters were established at a threshold of 7.5 pg for mouse T cells controlled by custom software coded in LabVIEW 2020 (National Instruments). Cells were sorted into 1.5 mL Eppendorf tubes, each filled with 35 uL of PBS+2%FBS, at a throughput of around 1,000 cells per hour. To maintain cell viability, all the T cell samples were preserved at 4 C before, during and after the sorting procedure.

### Bulk RNA-sequencing

For bulk RNA-sequencing, naive CD8+ T cells were sorted by mass into heavy (>7.5 pg) and light (<7.5 pg) T cell populations. At least 800 cells were sorted per condition. Sorted cell samples were brought to a volume of 250 uL with PBS and then 750 uL of Trizol (Invitrogen, 10296010) was subsequently added. Samples were then incubated for 5 minutes at room temperature. Following incubation, samples were transferred to 2 mL maXtract high density tubes (Qiagen, 129056) and 200 uL of chloroform was added. Samples were then vigorously mixed at room temperature for 2 minutes. To separate phases, samples were centrifuged for 5 minutes at 4 C and 12000xg. The aqueous phase was collected and mixed 1:1 with 70% ethanol. Following this step, the RNeasy Plus Micro Kit (Qiagen, 74034) was followed per manufacturer’s protocol starting at step 6. To briefly summarize the protocol, the RNA is collected in the RNeasy MinElute spin column, then washed with Buffer RW1, Buffer RPE, and 80% ethanol, and finally RNA is eluted in 14 uL of RNA-free water. After elution, RNA samples were stored at - 80 C until they were sequenced.

Samples were sent to MIT BioMicroCenter for library prep and subsequent RNA sequencing. Nucleic acid was treated with DNase (NEB), purified using RNA SPRI beads, and quantified using an Agilent FemtoPulse. cDNA was generated using Takara SMART-Seq® v4 Ultra® Low Input RNA Kit for Sequencing and Singular libraries prepared using Illumina NexteraXT with modified amplification oligonucleotides containing Singular S1/S2 anchors instead of Illumina P5/P7 (DNAscript Syntax). Libraries were sequenced on a Singular G4 sequencer using 150nt paired end reads.

Raw sequencing reads were aligned to the mmu10 reference genome using STAR, and transcripts were quantified using feature counts. Lowly expressed genes with fewer than 10 counts across more than 3 samples were filtered out. DESeq2 was used to normalize data and identify differentially expressed genes from the resulting count matrix. Genes with adjusted p values < 0.05 and absolute fold changes > 1.5 were considered significantly differentially expressed. The KEGGREST package was used to compile genes from relevant KEGG pathways for plotting with pheatmap. The set of mitochondrial genes was produced by selecting all genes beginning with the string “mt-”.

### Bulk T cell activation (>1 million T cells activated)

Mouse CD8+ T cells were activated with plate bound anti-CD3 (10 ug/mL, clone 145-2C11, BioXCell) and anti-CD28 (2 ug/mL, clone 37.51, BioXCell) in a tissue culture treated 24 well plate. Cells were seeded at a concentration of 1 million per well in T cell activation media. T cell activation media used was composed of RPMI (Invitrogen, 11875085) supplemented with 10% FBS (Gibco, 10438-026), 1% P/S (Gibco, 15140-122), 1x NEAA (Gibco, 11140-050), 1 mM sodium pyruvate (Gibco, 11360-070), 55 μM 2-mercaptoethanol (Gibco, 21985-023), 1x ITS (Gibco, 41400-045), and 10ng/mL IL-2 (StemCell, 78081). Cells were then maintained in the incubator at 37 C for 28 hours. At the 28 hour time point, cells were washed from the plate and stained with Ki67 and fixable viability dye for sorting of live cells immediately followed with fluorescent coupled SMR measurements.

### Low T cell count activation (<1000 T cells activated)

T cells sorted by buoyant mass into light and heavy subpopulations were activated and expanded for 6 days. T cells were activated with Dynabeads Mouse T-activator CD3/CD28 for T-cell expansion and activation (GIBCO, 11456D). Cells were seeded with beads at a concentration of 200 cells per well with a 1:8 cell:bead ratio in the InSphero Akura 96 spheroid microplate (InSphero, CS-PB15). Once T cells, beads, and media were all added to the wells, the whole plate was centrifuged for 6 minutes at 600xg to promote cell and bead contact. After centrifugation, the supernatant was removed and T cell activation media was added (RPMI supplemented with 10% FBS, 1% P/S, 1x NEAA, 1mM sodium pyruvate, 55 μM 2-mercaptoethanol, 1x ITS, and 10ng/mL IL-2). The plate was then stored in the incubator at 37 C for 6 days. Wells were imaged daily and on the 6th day, cells were stained with FC Block (101302), eBioscience Fixable Viability Dye (APC-Cy7, 65-0865-14), Ki67 (FITC, 11-5698-80), CD279 (PD-1) (Super Bright 600, 63-9985-80), CD366 (Tim3) (BV421, 119723), IFNg (APC, 17-7311-81), CD137 (4-1BB) (PE, 12-1371-81), CD62L (APC, 17-0621-81), and Granzyme B (Pacific Blue, 515407) for analysis via flow cytometry.

### Tumor conditioned media generation

KP-SIY cell line was grown at 37 °C and 5% CO_2_ in standard DMEM medium (Sigma, D5796-500ML) supplemented with 10% FBS (Gibco, 10438-026), 1% penicillin/streptomycin (GIBCO, 15140-122), and 2 ug/mL puromycin (Sigma, P8833-25mg). Media was removed from the cells after 7 days of culture (3 days after confluency) and filtered with a 0.2 μm filter to remove any remaining cells or debris. This media was stored at -80 C until the time of use. Media was thawed immediately before activation and mixed 1:9 with T cell activation media.

### T cell activation imaging

Over the course of the 6 day T cell activation, wells were visualized daily on the Nikon Eclipse TS100 using a 4x objective. Brightfield images were captured using the FLIR Blackfly S Monochrome Camera (Edmund Optics, BFS-U3-04S2M-CS).

### Flow cytometry

For flow cytometric analyses, cells were resuspended in staining buffer (Rockland, MB-086-0500) plus eBiosciences Fixable Viability Dye eFluor 780 and anti-CD16/CD32 and incubated at 4 C for 15 minutes. Cells were washed and then stained for surface proteins with fluorophore conjugated specific antibodies at 4 C for 30 min. If intracellular staining was required, eBioscience™ Intracellular Fixation & Permeabilization Buffer Set (Invitrogen, 88-8824-00) was used. Washed cells were resuspended in 100 uL of the fixation buffer and incubated in the dark at room temperature for 20 minutes. Cells were then washed twice in 1X Permabilization buffer. After completion of the washes, cells were resuspended in 100 uL of the permeabilization buffer with fluorophore conjugated specific antibodies and incubated for 20 minutes at room temperature in the dark. After a washing step, cells were immediately taken to flow cytometry for sorting.

### Amnis Imagestream measurement of cell shape and fluorescence

To quantitatively evaluate light vs heavy T cells, sorted T cells were stained and imaged on the Amnis Imagestream. Images were collected using the duo channel brightfield setting. Cell parameters of both light and heavy were analyzed using Amnis IDEAs. Raw data were output and plotted using GraphPad Prism. Fluorescent gatings were collected based on single color controls and fluorescent-minus-one gatings.

### Proteomic sample preparation

To prepare heavy and light CD8+ naïve T cell samples for MS, we adapted the OneTip protocol^24^ for use with SDB-RPS StageTips (CDS Analytical) and autosampler injection. SDB-RPS tips were first conditioned and equilibrated with 50 μL methanol followed by 50 uL 80% acetonitrile (ACN) with 0.1% formic acid (FA) and two volumes of 50 μL 0.1% FA. Centrifugation was performed at 1000 x *g* after each addition. SDB-RPS tips were pre-treated with 1 μL master mix (0.2% DDM, 100 mM triethylammonium bicarbonate buffer, 20 ng/μL trypsin (Promega), 10 ng/μL Lysyl Endopeptidase (Fujifilm Wako)) to passivate the SDB-RPS material. One thousand sorted heavy or light T cell suspensions from three independent mice (biological triplicate) in phosphate buffered saline (PBS) were loaded onto the tips and cells were concentrated on the SDB-RPS membrane by centrifugation at 300 x *g* until ∼2 μL remained. Two microliters of master mix were added to the 2 μL cell suspension and the tips were placed into microcentrifuge tubes with water, such that the level of the water reached ∼1 mm above the bottom of the SDB-RPS membrane. The tips and microcentrifuge tubes were then placed into a humidified ThermoMixer at 37°C for 4 hr for digestion. After digestion, the tips were removed and 50 μL 1% trifluoracetic acid was added, centrifuging at 1000 x *g* to bind peptides to the SDB-RPS material. Peptides were washed with 20 μL 0.1% FA followed by two volumes of 20 μL 80% ACN with 0.1% FA. Peptides were eluted directly into glass autosampler vials with 20 μL 5% ammonium hydroxide in 80% ACN. Eluted peptides were dried using a SpeedVac before reconstitution in 1 μL 3% ACN, 0.1% FA with 0.015% DDM and brief sonication to promote resuspension.

A bulk collection of naïve T cells was processed for spectral library generation using S-Trap spin columns (Protifi) according to the vendor’s instructions, with the exception that reduction and alkylation were not performed. Half of the resulting peptides were fractionated offline using a high-pH reversed-phase fractionation kit (Pierce), eluting in six fractions (7.5% ACN with 0.093% triethylamine (Et_3_N), 10% ACN with 0.090% Et_3_N, 12.5% ACN with 0.088% Et_3_N, 15% ACN with 0.085% Et_3_N, 17.5% ACN with 0.083% Et_3_N, and 50% ACN with 0.05% Et_3_N).

### Liquid Chromatography-Tandem Mass Spectrometry (LC-MS/MS) Analysis

LC–MS/MS analyses were performed on a Bruker timsTOF Ultra mass spectrometer coupled to a Bruker nanoElute2 LC system. The LC system was equipped with a PepSep 25 cm x 75 μm inner diameter x 1.5 μm C_18_ particle analytical column, operated at a 200 nL/min flow rate and interfaced with a 10 μm PepSep emitter and Bruker Captive Spray Ultra source. The column was heated to 50°C using a Bruker column toaster.

The full sample of heavy and light CD8+ T cells was injected and separated using a 30 min linear gradient from 3–34% mobile phase B (ACN with 0.1% FA). Mobile phase A was water with 0.1% FA. MS acquisition was performed in diaPASEF mode, with an optimized 16 x 3 diaPASEF window scheme that was generated using the py_diAID tool^25^ before manual adjustment. Our optimized window scheme considered precursors ranging from 1/K0 = 0.65 to 1.46 Vs cm^−2^ and 302 to 1298 m/z. The trapped ion mobility spectrometry (TIMS) ramp time was set to 100 ms and low sensitivity mode was enabled. Collision energy was scaled as a function of ion mobility starting from 20.00 eV at 0.60 Vs cm^−2^ to 59.00 eV at 1.60 Vs cm^−2^.

For spectral library generation, 150 ng peptide from bulk naïve T cell was injected and analyzed using diaPASEF, as above, whereas for fractionated bulk naïve T cell sample, 150 ng peptide was injected per fraction and separated using a 180 min linear gradient from 2–35% mobile phase B. MS acquisition was performed in ddaPASEF mode, with one TIMS-MS survey scan and 10 PASEF MS/MS scans acquired per acquisition cycle. Precursor ions in the mobility range of 1/K0 = 0.7 to 1.45 Vs cm^−2^ and 302 to 1298 m/z were considered for MS/MS analysis using a 2 m/z isolation window at 700 m/z and 3 m/z at 800 m/z. Precursor ions were selected with an intensity threshold of 500 arbitrary units (a.u.) and dynamically excluded for 40 sec or until reaching a 20,000 a.u. target intensity. The TIMS ramp time was set to 100 ms and the collision energy was scaled as in the diaPASEF method.

### MS Data Analysis

To generate the ddaPASEF spectral library, ddaPASEF raw files were searched using FragPipe (version 22.0) against a UniProt mouse database (UP000000589, downloaded Oct 2024). DDA+ mode was enabled. Tryptic peptides ranging from 7–50 amino acids and 500–5,000 m/z with a maximum of two miscleavages were considered in MSFragger^26,27^. Methionine oxidation and N-terminal acetylation were set as variable modifications and the precursor and fragment mass tolerances were set to 10 and 20 ppm, respectively. MSBooster^28^ was enabled with the DIA-NN retention time prediction and Prosit 2023 Intensity timsTOF spectra prediction models selected. Percolator^29^ was used for peptide sequence match validation with the minimum probability threshold set to 0.5. The false discovery rate (FDR) was capped at 1%.

diaPASEF raw files were first searched using FragPipe to generate a DIA-based spectral library by enabling diaTracer diaPASEF spectrum deconvolution^30^. The resulting diaPASEF library was then combined with the ddaPASEF library using DIA-NN^31^ (version 1.9.2) to generate a hybrid diaPASEF/ddaPASEF spectral library. Finally, diaPASEF raw files were searched against the hybrid library for quantification using DIA-NN^32^. The maximum number of miscleavages was capped at 1, N-terminal M excision was enabled, and C carbamidomethylation was disabled. Precursor ions ranging from 7–30 amino acids, 1–4 *z*, and 300–1800 m/z and fragment ions from 200–1800 m/z were considered. The precursor FDR was set to 1% and the MS1 and MS2 mass accuracies were set to 15 ppm each. The quantification strategy was set to QuantUMS (high precision), and cross-run normalization was disabled.

DIA-NN output matrices were imported into FragPipe Analyst^33^ for further analysis. Missing values were imputed using minimum value imputation before applying a local and tail area-based FDR correction. Proteins with a log_2_ fold change value of ≥ 0.58 or ≤ –0.58 and an adjusted p-value of <0.05 were considered differently abundant between heavy and light CD8+ T cells. Differently abundant proteins were submitted for pathway enrichment and cellular component analyses using STRING^34^. Tissue-specific functional module detection using HumanBase (Flatiron Institute – Simons Foundation)^35,36^ was performed with a human T lymphocyte background upon converting mouse gene names to the human equivalent. The StringApp Plugin within Cytoscape^37^ (version 3.10.3) was used for sub-clustering based on gene ontology biological process, cellular component, molecular function and Reactome pathway classification, as well as to acquire protein-protein interactions, with edges set to medium confidence and all interaction sources active (textmining, experiments, databased, co-expression, neighborhood, gene fusion, and co-occurrence)^34,38^. Enrichment signal was defined as the weighted harmonic between the observed/expected ratio and the log(FDR).

## Supporting information

All extended data

## Statistical analysis

The results are shown as mean ± standard error of the mean (SEM). To determine the statistical significance of the differences between the experimental groups two-tailed unpaired or paired Student’s t tests, 2-way ANOVA tests were performed using the Prism 10 software (GraphPad), as indicated in each figure. Sample sizes were based on experience and experimental complexity, but no methods were used to determine normal distribution of the samples. Differences reached significance with p values < 0.05 (p values noted in figures). The figure captions contain the number of independent experiments or mice per group that were used in the respective experiments.

## Reporting Summary

Further information on research design is available in the Nature Research Reporting Summary linked to this article.

## Data availability

The authors declare that all data supporting the findings of this study are available within the paper and its Supplementary Information. Source data for the primary-sample figures are available. MS data are deposited in the ProteomeXchange Consortium using the MassIVE repository with the project accession of PXD058965. This can be accessed at: https://massive.ucsd.edu/ProteoSAFe/private-dataset.jsp?task=ec3673edda5e4b3294d8a77b6897bc0d with the username “MSV000096684_reviewer” and the password “BuoyantMass”.

## Code availability

Custom SMR data processing and sequencing analysis code is available in the supplemental information.

## Acknowledgements

We thank the Brigham and Women’s Hospital Specimen Bank for providing human donor PBMC collar samples, and for Dr. Lydie Debaize, Dr. Mark Murakami, coordination. We thank Dr. Leyuan Ma, Dr. Kewen Lei and Dr. Darrell Irvine for helpful discussion in T cell activation, flow cytometry, and mouse work. We thank Kevin Yang for discussion and for helping the schematics in the figures. We thank the laboratory of Tyler Jacks for providing C57BL/6; 129/Sv mixed background mice. Some schematics presented in this work were created with BioRender.com.

## Author contributions

J.Y, Y.Z., and S.R.M. conceptualized the study on discovering and characterizing light and heavy T cells; R.J.K. and G.M.B. conceptualized the clinical study. Y.Z. designed and built the sorting device. For in vitro experiments, J.Y, S.M.D, T.U. and Y.Z designed and carried out the experiments. For in vivo experiments, T.D., S.M.D., J.Y., Y.Z designed and carried out the experiments. G.L.Y. and J.Y. analyzed the RNA-sequencing data, Y.Z assisted the analysis. S.C., A.L. and T.S. coordinated the clinical study, R.L., M.V., K.S., and R.A. performed sample processing and data collection under the supervision of R.J.K. and M.S. S.O., N.L.C., and R.J.K. performed the data analysis for the clinical specimens. J.Y., Y.Z., S.M.D and S.R.M. wrote the manuscript with contributions from R.J.K. All authors reviewed and approved the manuscript. S.R.M. provided overall project leadership.

