## Supplementary material for "Buoyant mass reveals distinct T cell states predictive of checkpoint response": All extended data

### Extended Data Table of Contents

**Extended Data Note 1: Relationship between cellular buoyant mass and dry mass** 2

**Extended Data Fig.1 | Isolating CD8+ T cells using flow cytometry.** 3

**Extended Data Fig.2 | Single cell sequencing of naive T cells before and after SMR.** 4

**Extended Data Fig.3 | FSC/SSC and Amnis Imagestream analysis of heavy and light T cells.** 5

**Extended Data Fig.4 | Mass-based SMR single cell sorting.** 6

**Extended Data Fig.5 | Bulk RNA sequencing of light and heavy T cells.** 8

**Extended Data Fig.6 | Gene pathway analysis on bulk RNA sequencing of light and heavy T cells.** 9

**Extended Data Fig.7 | Proteomic analysis of sorted heavy and light T cells.** 10

**Extended Data Fig.8 | CD3/CD28 Dynabeads T cell activation in normal and tumor conditioned T cell activation media.** 12

**Extended Data Fig.9 | Light and heavy activated T cells Amnis Imagestream gating.** 14

**Extended Data Fig.10 | Buoyant mass distributions of peripheral T cells from melanoma patients.** 15

**Extended Data Fig.11 | Additional analysis of single-cell biophysical measurements and their association with ICB treatment response.** 16

**Extended Data Note 1: Relationship between cellular buoyant mass and dry mass**

Cell buoyant mass, $m_b$, includes a component from the dry material and another from the intracellular water content as given by

$$m_b = m_{dry}(1 - \frac{\rho_{fluid}}{\rho_{dry}}) + V_{iw}(\rho_{iw} - \rho_{fluid}) \quad (1)$$

where $\rho_{fluid}$ is the fluid density, $m_{dry}$ and $\rho_{dry}$ are the mass and density of the cell's dry materials, and $V_{iw}$ and $\rho_{iw}$ are the volume and density of the exchangeable water content. When cells are in media where the density is close to that of water ($\rho_{iw} \approx \rho_{fluid}$), the buoyant mass depends on only dry mass and dry density

$$m_b = m_{dry}(1 - \frac{\rho_{fluid}}{\rho_{dry}}) \quad (2)$$

In our measurements, cells were suspended in 1X PBS ($\rho_{fluid} = 1.005\ g \cdot cm^{-3}$). Theoretically, when the dry density in equation (2) is constant across both light and heavy populations ($\rho_{dry} \approx 1.4\ g \cdot cm^{-3}$ as previously reported[1]), the buoyant mass is linearly correlated with the dry mass with a constant slope $k = 1/(1 - \frac{\rho_{fluid}}{\rho_{dry}}) \approx 3.5$. Based on our measurements, linear regression between buoyant mass and dry mass of the heavy population is consistent with this theoretical estimate while the light population does not match as well (**Extended Data Note 1 Fig.**). These calculations indicate that light and heavy T cells have different dry densities. Moreover, this observation appears in both human and mouse samples.

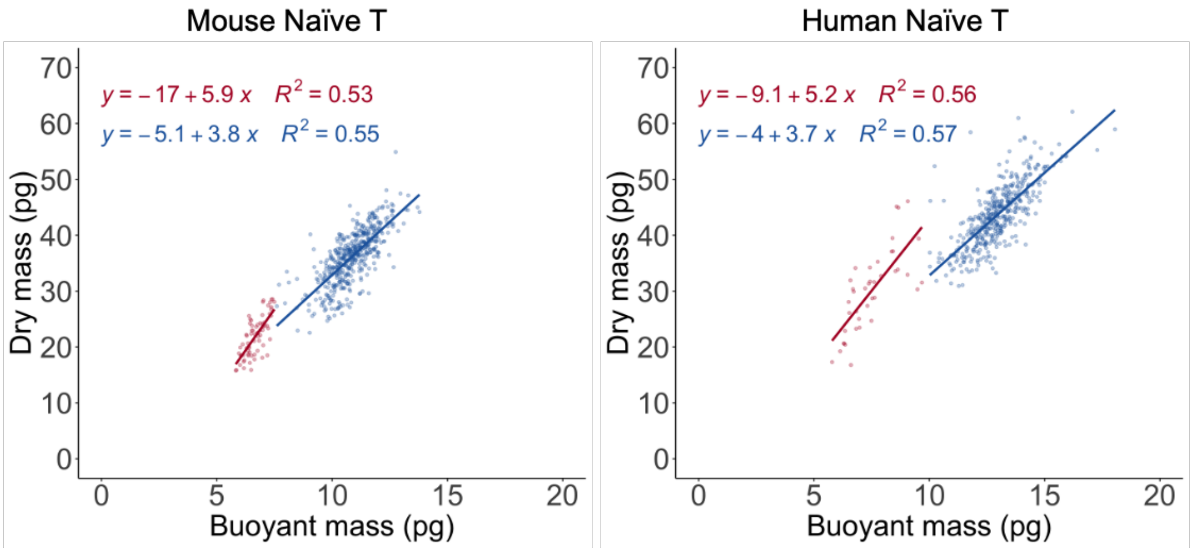


**Extended Data Note 1 Fig. | Relationship between cellular buoyant mass and dry mass**. Buoyant mass and dry mass of mouse and human naïve T cells. Light cells are represented in red while heavy ones in blue. Lines in the graphs represent the linear regression between buoyant mass and dry mass of each population.

**a.**

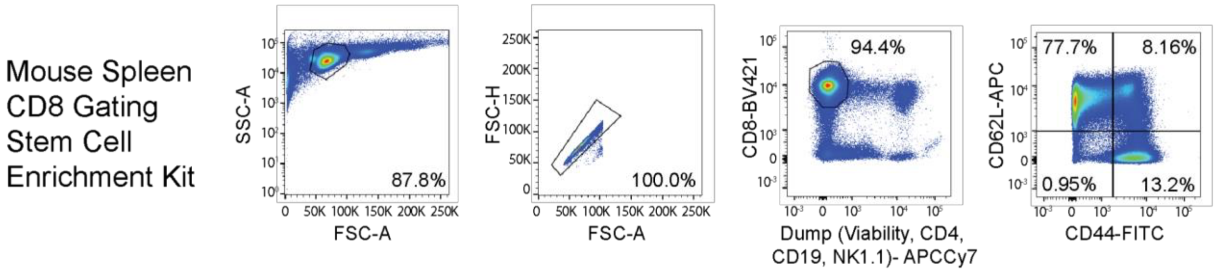


**Extended Data Fig.1 | Isolating CD8+ T cells using flow cytometry.**
**a,** Representative gating strategy to identify and sort naive (CD62L+, CD44-), central memory (CD62L+, CD44-), and effector memory (CD62L-, CD44+) CD8+ T cell subpopulations from healthy mouse spleen samples.

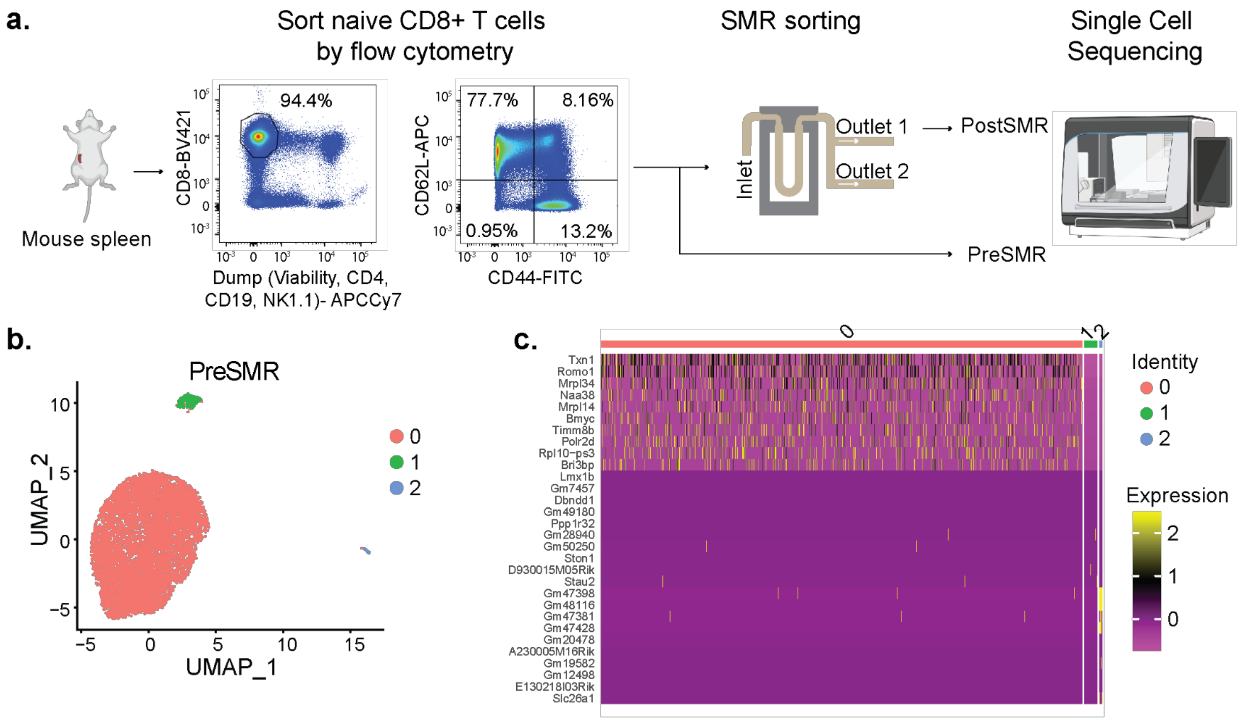


d. PreSMR Number of contaminating cells per cluster

| Cluster | B Cell | NK Cell | Monocyte | Neutrophil |
|---|---|---|---|---|
| 0 | 6 | 11 | 0 | 0 |
| 1 | 0 | 0 | 0 | 0 |
| 2 | 0 | 0 | 0 | 0 |

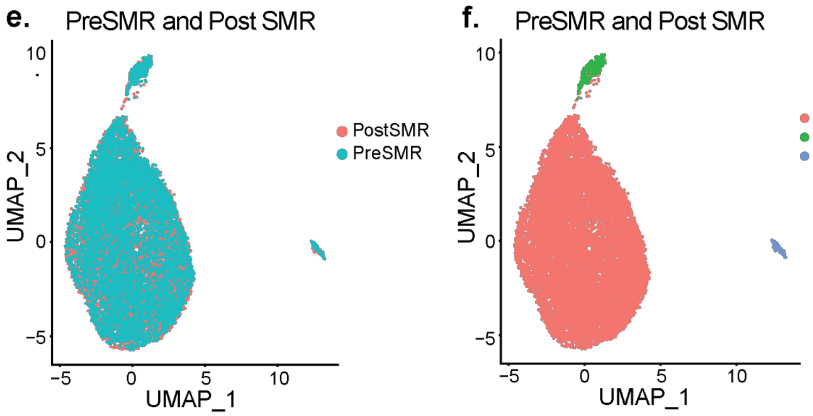


g. PostSMR cell counts per cluster

| Cluster | PreSMR | PostSMR |
|---|---|---|
| 0 | 7567 | 7148 |
| 1 | 240 | 120 |
| 2 | 81 | 122 |

**Extended Data Fig.2 | Single cell sequencing of naive T cells before and after SMR.**
**a,** Schematic of the workflow for preparing Naive T cells before and after SMR sorting for single cell sequencing. **b,** UMAP of flow sorted naive T cells (preSMR, 7627 cells in total) defined by 3 clusters. **c,** Heatmap of differentially expressed genes between 3 naive T cell (naive T cells preSMR) UMAP clusters. **d,** Number of potential immune cell contaminates per cluster based on expression of 3 or more defined surface markers (**Supplemental Table 4**). **e,** UMAP of naive T cells collected after flow sorting (preSMR) and being flown through the SMR (postSMR) colored by sample. **f,** UMAP of preSMR and postSMR naive T cells defined and colored by 3 clusters. **g,** Number of cells from each sample per cluster.

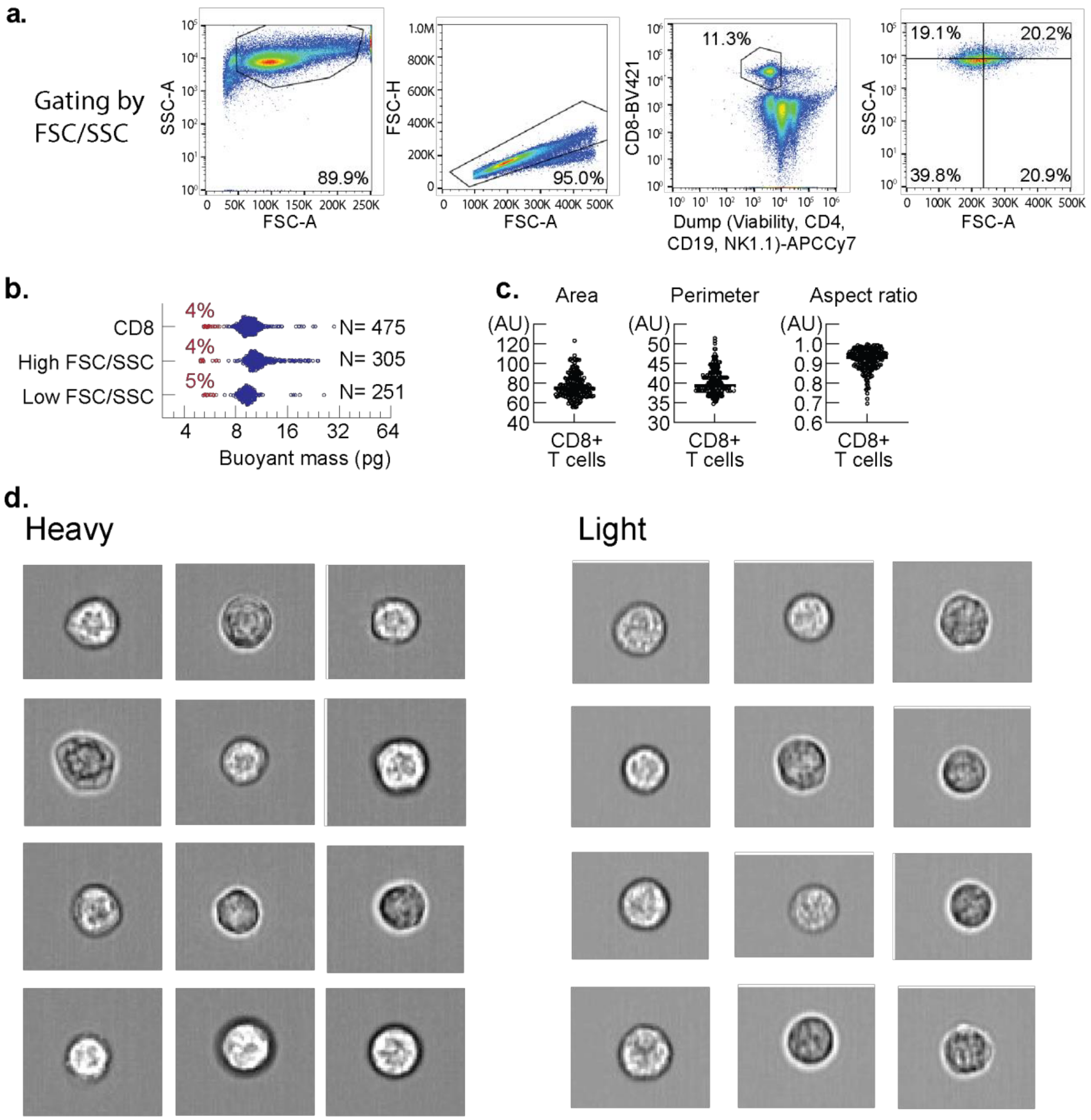


**Extended Data Fig.3 | FSC/SSC and Amnis Imagestream analysis of heavy and light T cells.**
**a,** Representative gating strategy to sort CD8 T cells by light scattering. Samples from Q2 (i.e. High SSC-A high FSC-A) and Q4 (low SSC-A low FSC-A) were collected and measured with the SMR. **b,** Mass distribution of CD8+ T cells and CD8+ T cell populations sorted by FSC/SSC. **c,** Measurements of total cell area, perimeter, and aspect ratio of mouse CD8+ T cells on the Amnis® Imaging Flow Cytometer. **d,** Representative brightfield image from the Amnis Imagestream of heavy (left) and light (right) mouse spleen CD8+ T cells.

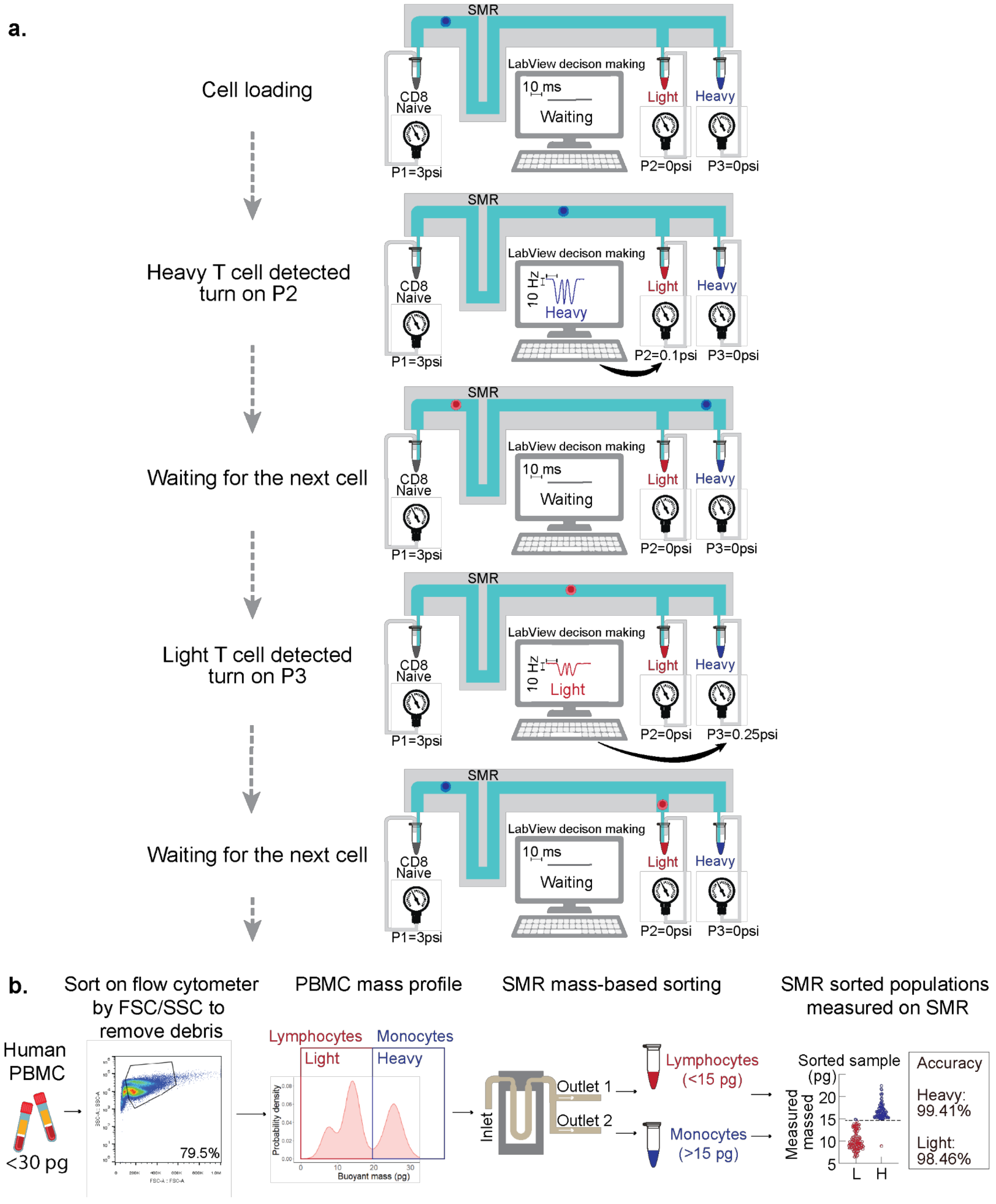


**Extended Data Fig.4 | Mass-based SMR single cell sorting.**
**a,** T cells (naive or CD8) are loaded onto the system with positive pressure from regulator 1 (P1). As the cells flow through the cantilever on SMR, the mass of the cell is recorded as frequency shift (Hz). A decision is made in LabVIEW based on the frequency shift and pre-set thresholds to control the subsequent regulators P2 and P3. For example, a mouse T cell heavier than 12.5 Hz is considered heavy

and will trigger pressure regulator P2 to direct the cell into the heavy collector. On the contrary, a light cell will trigger regulator P3 to direct the cells into the light collector. The system allows only one cell in the fluidic path at any given time to ensure sorting accuracy. The process repeats for each cell automatically. Cells are sorted at a rate of about 1000 cells per hour. **b,** A PBMC sample was used to test sorting accuracy. Lymphocytes and monocytes were collected from a human PBMC sample through FSC-SSC sorting on flow cytometry. Monocytes are distinctly heavier than lymphocytes. We sorted monocytes and lymphocytes based on mass. The sorted samples were then measured on SMR to evaluate the sorting accuracy. Accuracy is defined as the percentage of cells in the light (or heavy) compartment that is measured to be light. The heavy compartment has an accuracy of 99.41% and the light 98.46%.

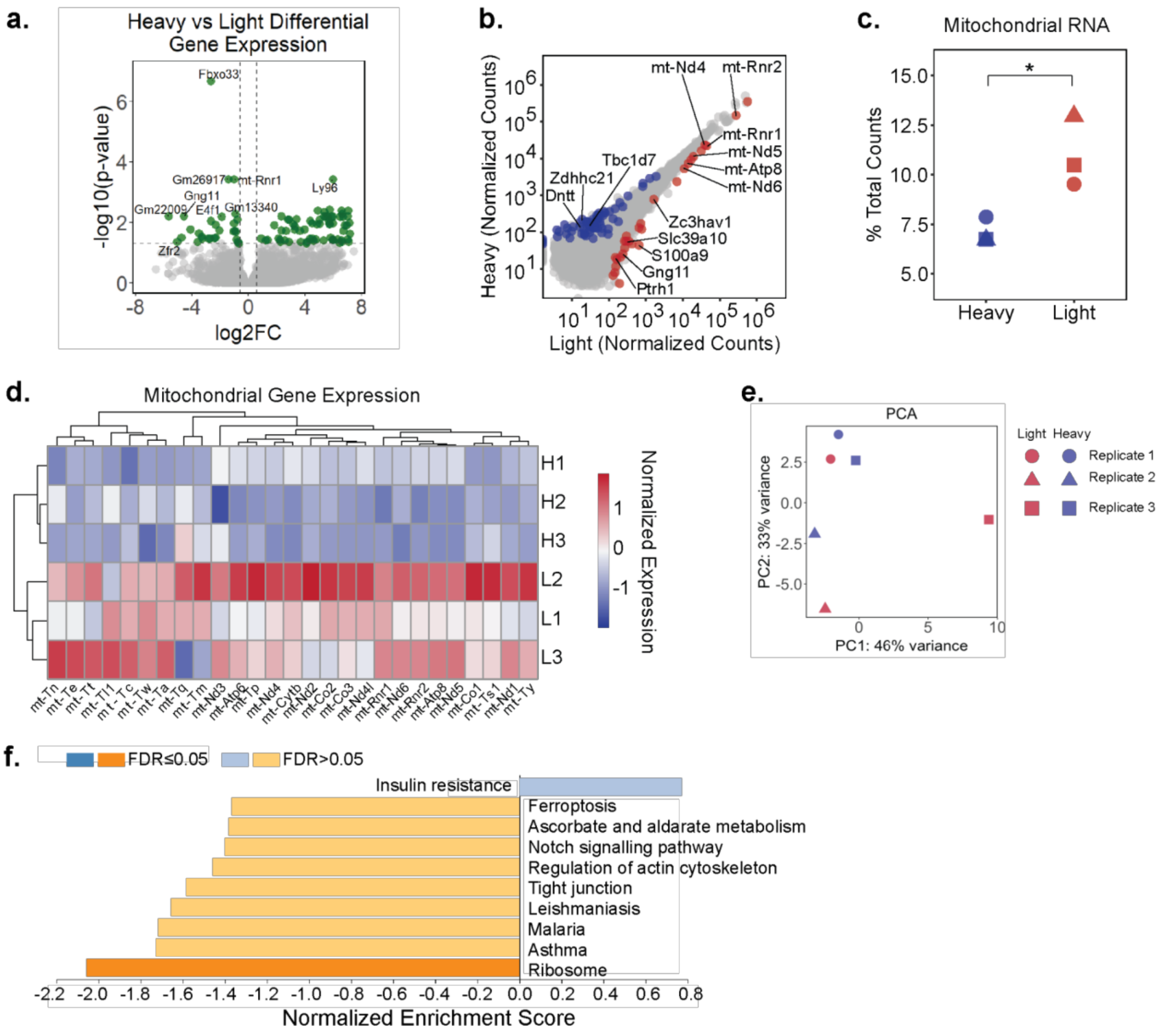


**Extended Data Fig.5 | Bulk RNA sequencing of light and heavy T cells.**
**a,** Volcano plot of heavy vs. light differential gene expression. Heavy cells were used as reference and fold change numbers represent light cells mRNA expression levels compared to heavy. b, Normalized gene counts for heavy and light T cells. Differentially expressed genes are colored (adjusted p-value ≤ 0.05 and absolute fold change ≥ 1.5). Blue denotes upregulation in heavy cells, and red denotes upregulation in light cells. **c**, Comparison of the percentage of total mitochondrial RNA (unpaired t-test, p-value = 0.049). **d**, Heatmap of normalized counts for all expressed mitochondrial genes. Hierarchical clustering (complete, Euclidean distance) is performed on samples and genes. **e.** Principal component analysis on heavy and light T cells. Shapes depict different replicates and color depicts buoyant mass. **f.** KEGG gene set enrichment analysis was performed to compare light and heavy T cells. Heavy cells were used as reference and enrichment scores represent light cells mRNA expression levels compared to heavy.

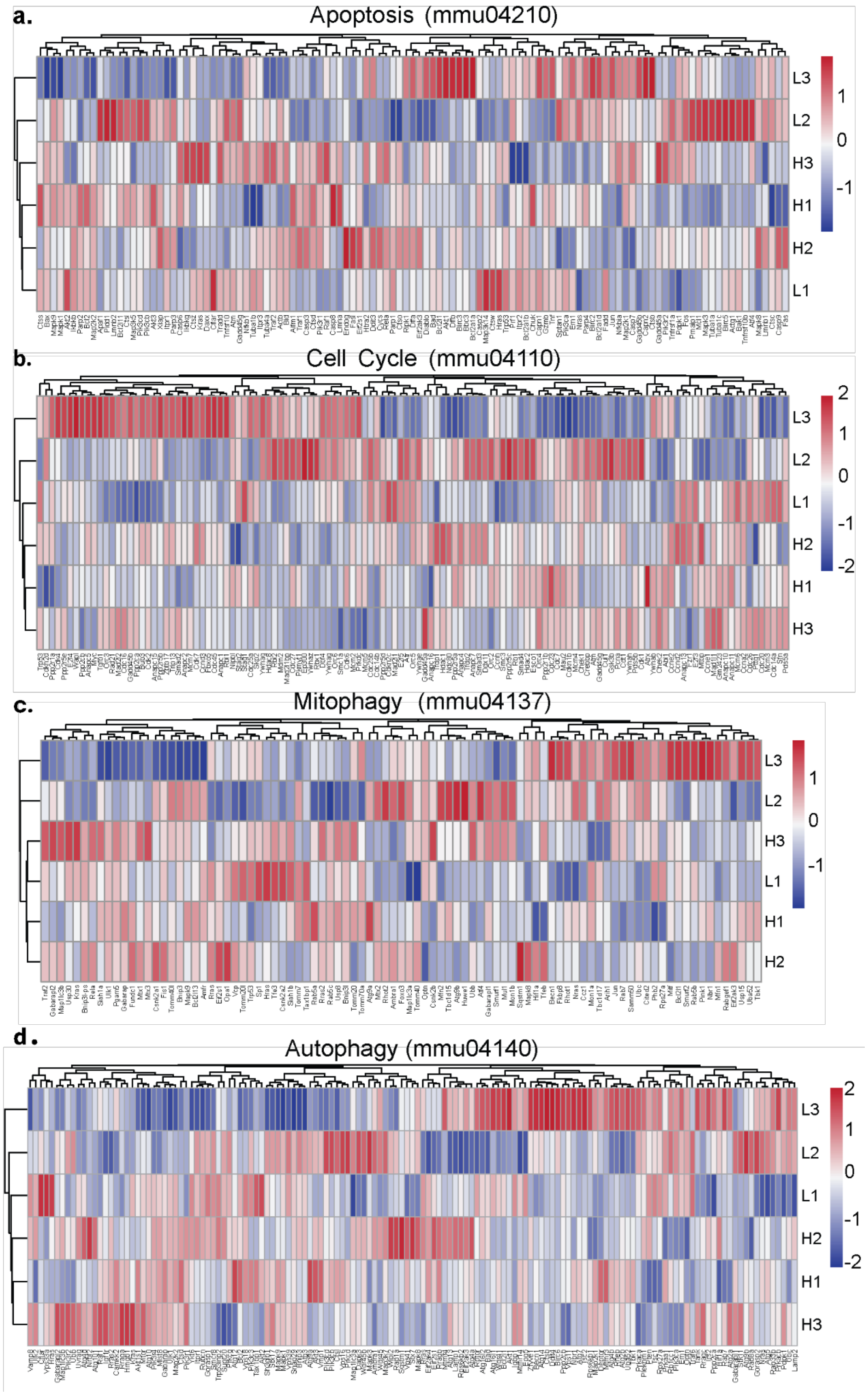


**Extended Data Fig.6 | Gene pathway analysis on bulk RNA sequencing of light and heavy T cells.** Heatmap visualization of genes related to pathways of interest. Pathways analyzed are **a,** apoptosis, **b,** cell cycle, **c,** mitophagy, and **d,** autophagy.

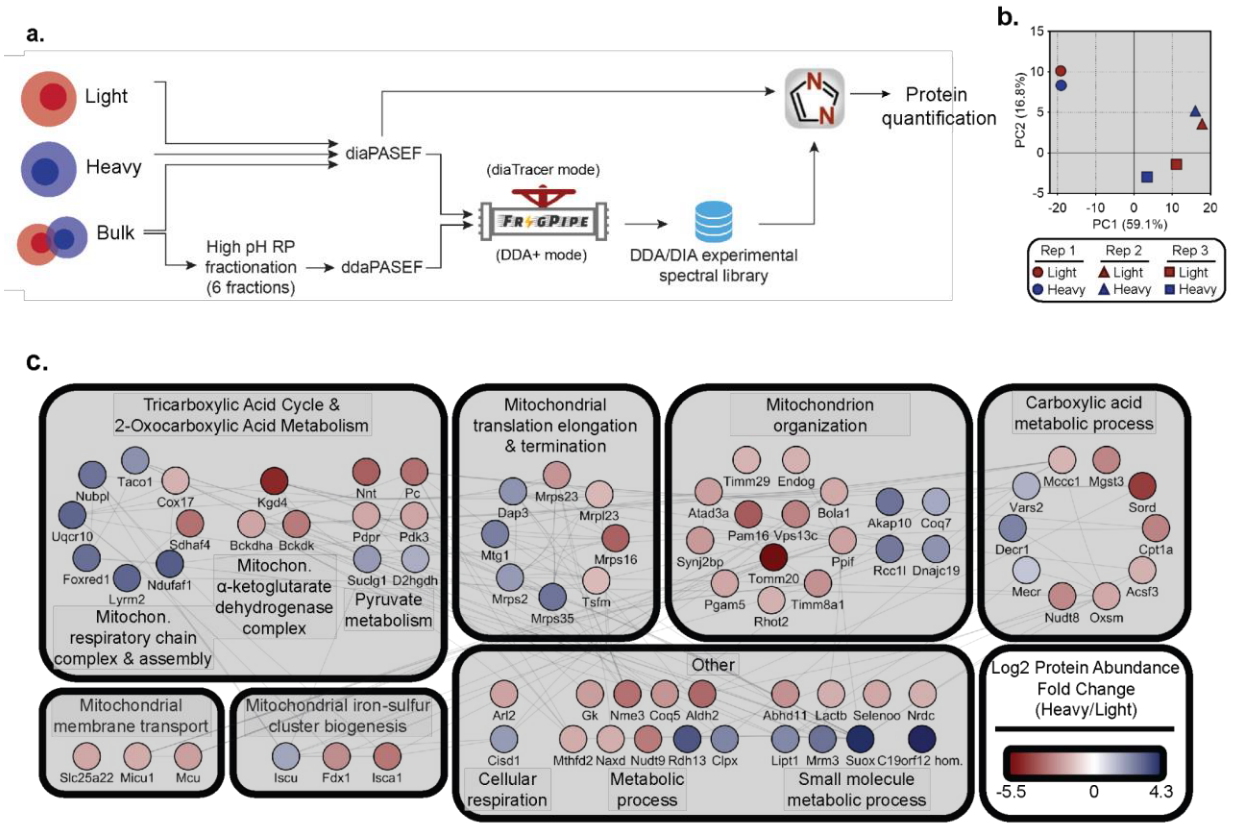


**Extended Data Fig.7 | Proteomic analysis of sorted heavy and light T cells.**

**a,** Schematic of the mass spectrometry analysis workflow for heavy and light T cell proteomic samples. Heavy and light cells were analyzed on the mass spectrometer using parallel accumulation serial fragmentation in data-independent acquisition (diaPASEF)[2] mode. Bulk naïve T cells were analyzed in diaPASEF mode, and additionally fractionated by high pH reversed-phased (RP) fractionation for analysis in data-independent acquisition (dda)-PASEF (ddaPASEF) mode. For enhanced protein identification, a hybrid DDA/DIA experimental mass spectral library was generated in FragPipe[3]and DIA-NN6[4]. Heavy and light proteomic samples were further quantified by searching the raw mass spectrometry data against the hybrid spectral library using DIA-NN7[5]. **b,** Principal components analysis of heavy and light T cell proteomic samples from 3 mice. **c,** Combined protein-protein interaction network of mitochondrial proteins determined from **(Figure 4c,d),** containing those enriched in light cells (adjusted p-value ≤ 0.05 and log2 fold change ≤ -0.58) and those enriched in heavy cells (adjusted p-value ≤ 0.05 and log2 fold change ≥ 0.58). Protein nodes indicate the relative abundance as a ratio of heavy over light T cell samples. The STRING database was used to sub-cluster proteins based on pathway enrichment and for determining functional protein interactions (edges in gray).

a.

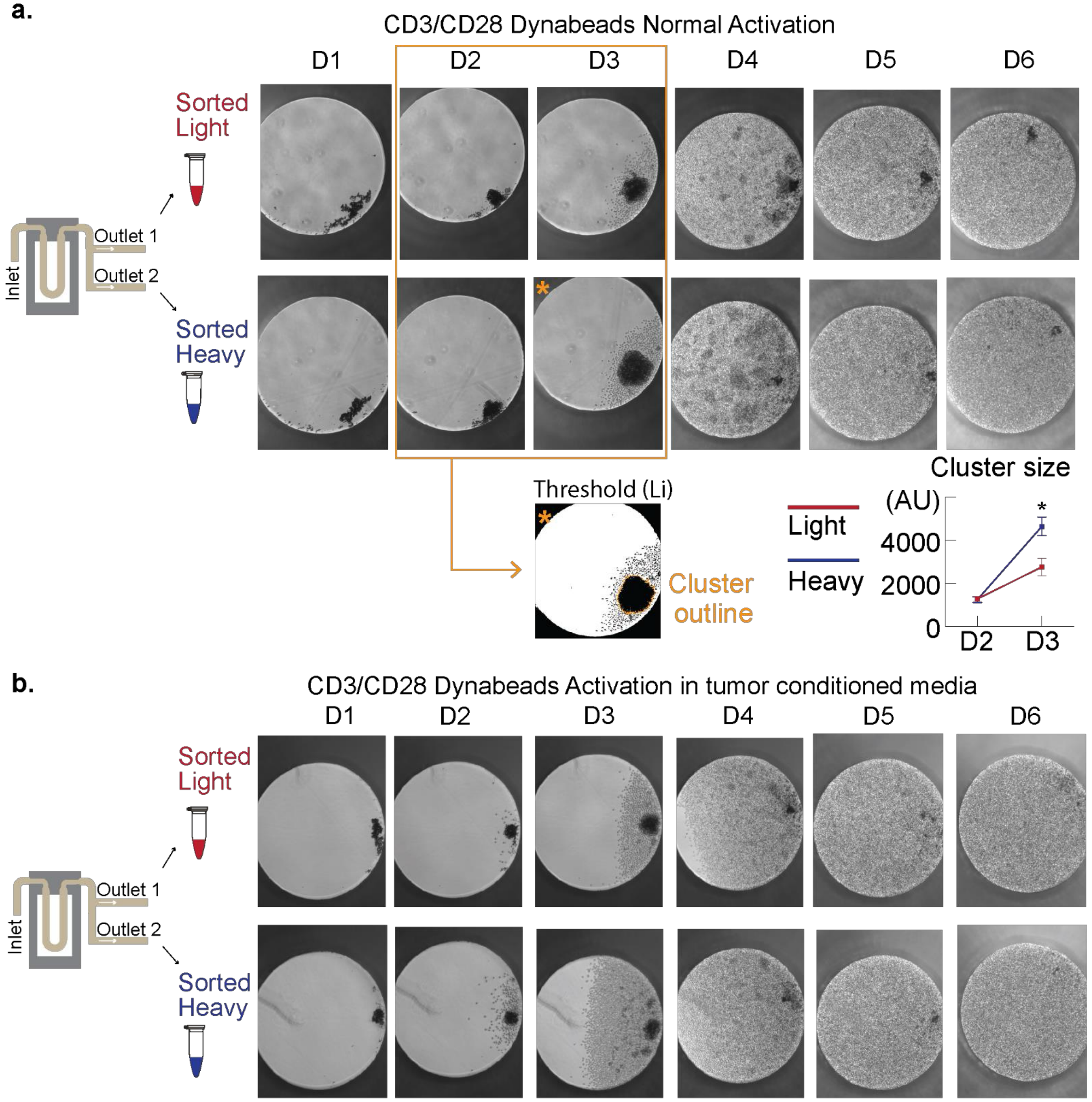


c.

Heavy and Light Activation D6 SMR measurement

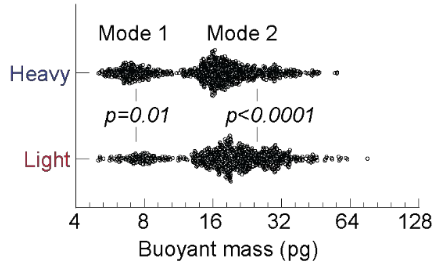


| | Heavy | Light |
|---|---|---|
| Mode 1 (pg) | 7.14 | 7.84 |
| Mode 2 (pg) | 16.53 | 18.28 |

**Extended Data Fig.8 | CD3/CD28 Dynabeads T cell activation in normal and tumor conditioned T cell activation media.**

**a,** T cells sorted by mass were activated for 6 days and imaged daily. A cluster starts to form on day 1. Importantly there is no proliferation so the cluster is formed by T cells migrating and carrying the beads together (data not shown). The cluster grows in the following days, due to both enlargement of T cells during activation and proliferation. Images on day 5 and 6 show that the wells are confluent with T cells. Analysis of cluster size was performed on days 2 and 3. Li thresholding in ImageJ is used to identify the cluster so that cluster area can be calculated. An example of the cluster outline is shown in orange and the image used for analysis is indicated with a star. **b,** T cells sorted by mass were activated in the presence of 10% tumor conditioned media for 6 days and imaged daily. Similar to activation without tumor conditioned media, clusters start to form on day 1. Cluster grows in the following days, due to both enlargement of T cells during activation and proliferation. Images on day 5 and 6 are confluent with T cells. **c,** CD8+ T cells sorted by mass into light and heavy bins were independently activated and subsequently measured on the SMR after 6 days. Modes of each mass distribution were calculated using the package multimode in R[6] and tabulated. Statistical analysis was performed using two-tailed unpaired t-tests with Welch's correction.

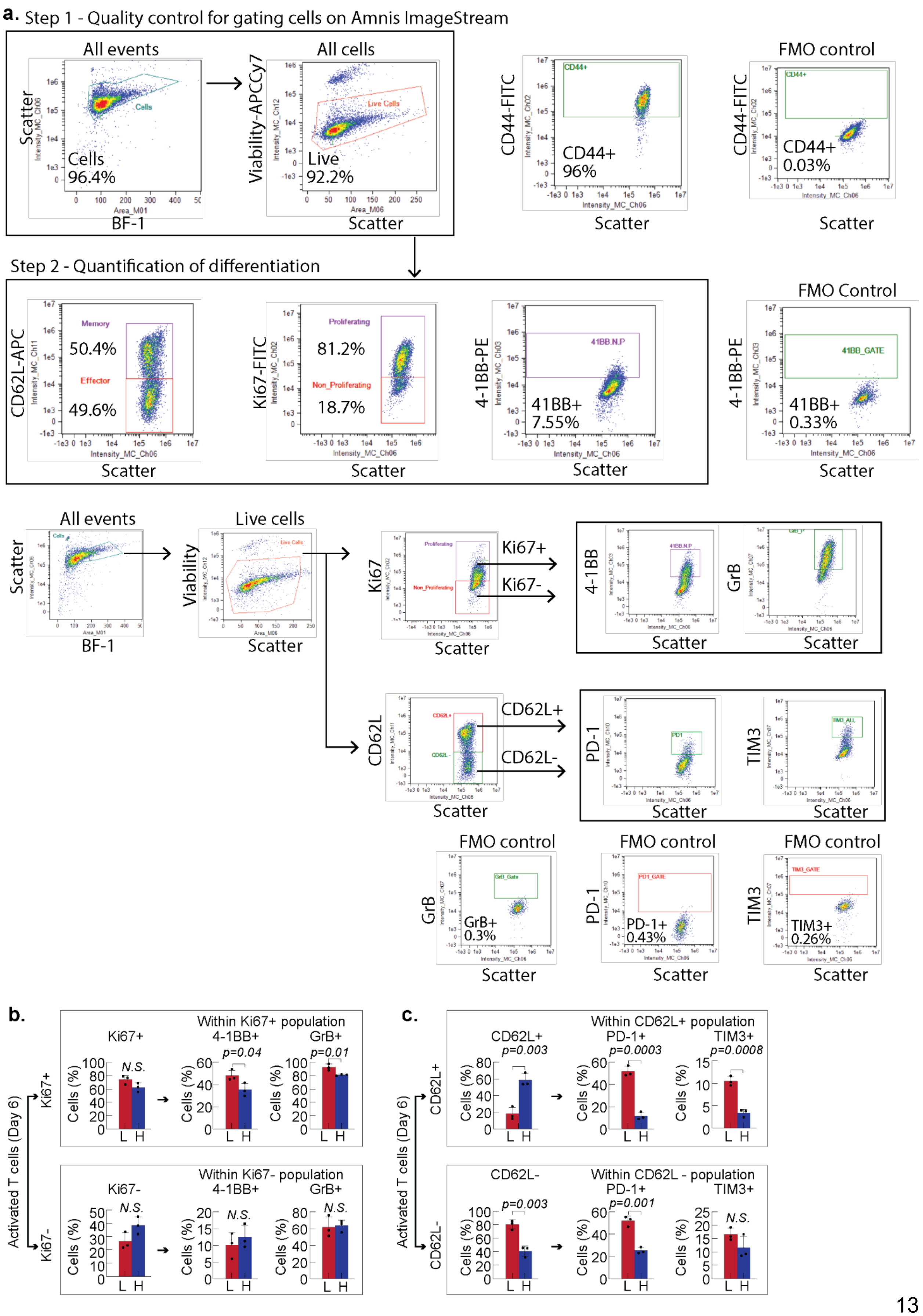

a. Step 1 - Quality control for gating cells on Amnis ImageStream
All events
All cells
FMO control
Scatter
Cells 96.4%
BF-1
Viability-APCCy7
Live 92.2%
Scatter
CD44-FITC
CD44+ 96%
Scatter
CD44-FITC
CD44+ 0.03%
Scatter
Step 2 - Quantification of differentiation
FMO Control
CD62L-APC
Memory
50.4%
Effector
49.6%
Scatter
Ki67-FITC
Proliferating
81.2%
Non_Proliferating
18.7%
Scatter
4-1BB-PE
41BB+ 7.55%
Scatter
4-1BB-PE
41BB+ 0.33%
Scatter
All events
Live cells
Scatter
BF-1
Viability
Scatter
Ki67
Ki67+
Ki67-
Scatter
4-1BB
GrB
Scatter
Scatter
CD62L
CD62L+
CD62L-
Scatter
PD-1
TIM3
Scatter
Scatter
FMO control
FMO control
FMO control
GrB
GrB+ 0.3%
Scatter
PD-1
PD-1+ 0.43%
Scatter
TIM3
TIM3+ 0.26%
Scatter
b.
Within Ki67+ population
Ki67+
4-1BB+
GrB+
N.S.
p=0.04
p=0.01
Within Ki67- population
Ki67-
4-1BB+
GrB+
N.S.
N.S.
N.S.
Activated T cells (Day 6)
Ki67+
Ki67-
Cells (%)
L H
c.
Within CD62L+ population
CD62L+
PD-1+
TIM3+
p=0.003
p=0.0003
p=0.0008
Within CD62L - population
CD62L-
PD-1+
TIM3+
p=0.003
p=0.001
N.S.
Activated T cells (Day 6)
CD62L+
CD62L-
Cells (%)
L H

**Extended Data Fig.9 | Light and heavy activated T cells Amnis Imagestream gating.**
**a,** Representative gating strategy to identify differentiated CD8+ T cell subtypes from activated light and heavy T cells using the Amnis Imagestream. **b,** Amnis Imagestream analysis of Ki67 expression and expression of granzyme B and 4-1BB within both Ki67+ and Ki67- populations on day 6 of activation. **c**, Amnis Imagestream analysis of CD62L expression and expression of T cell exhaustion markers TIM3 and PD-1 within the CD62L+ and CD62L- populations on day 6 of activation. (**b**) & (**c**) Data are shown as mean ± s.e.m., representative of three independent experiments. Statistical analysis was performed using two-tailed unpaired t-tests with Welch's correction.

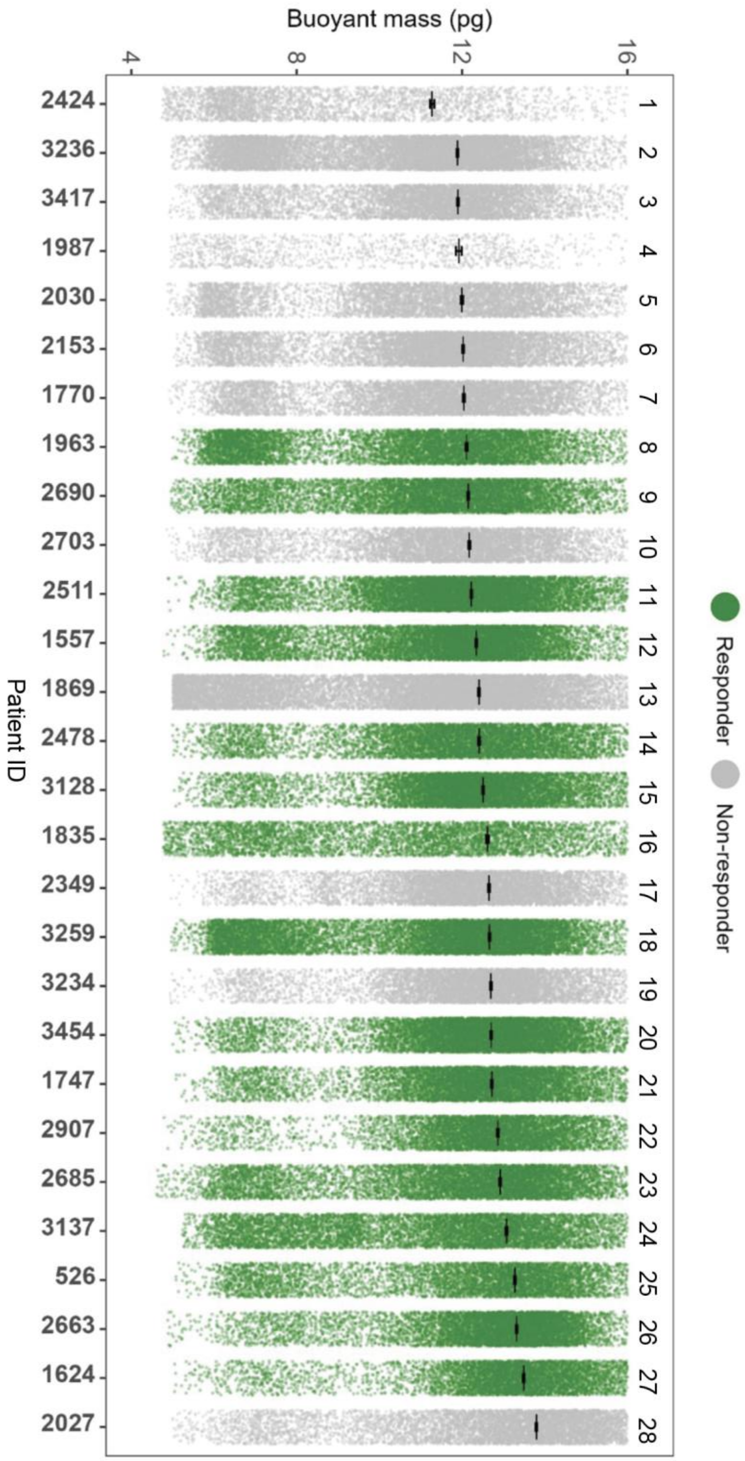


**Extended Data Fig.10 | Buoyant mass distributions of peripheral T cells from melanoma patients.** Buoyant mass distributions of pretreatment peripheral blood T cells from 28 patients, ordered by heavy mean and colored by response to ICB therapy.

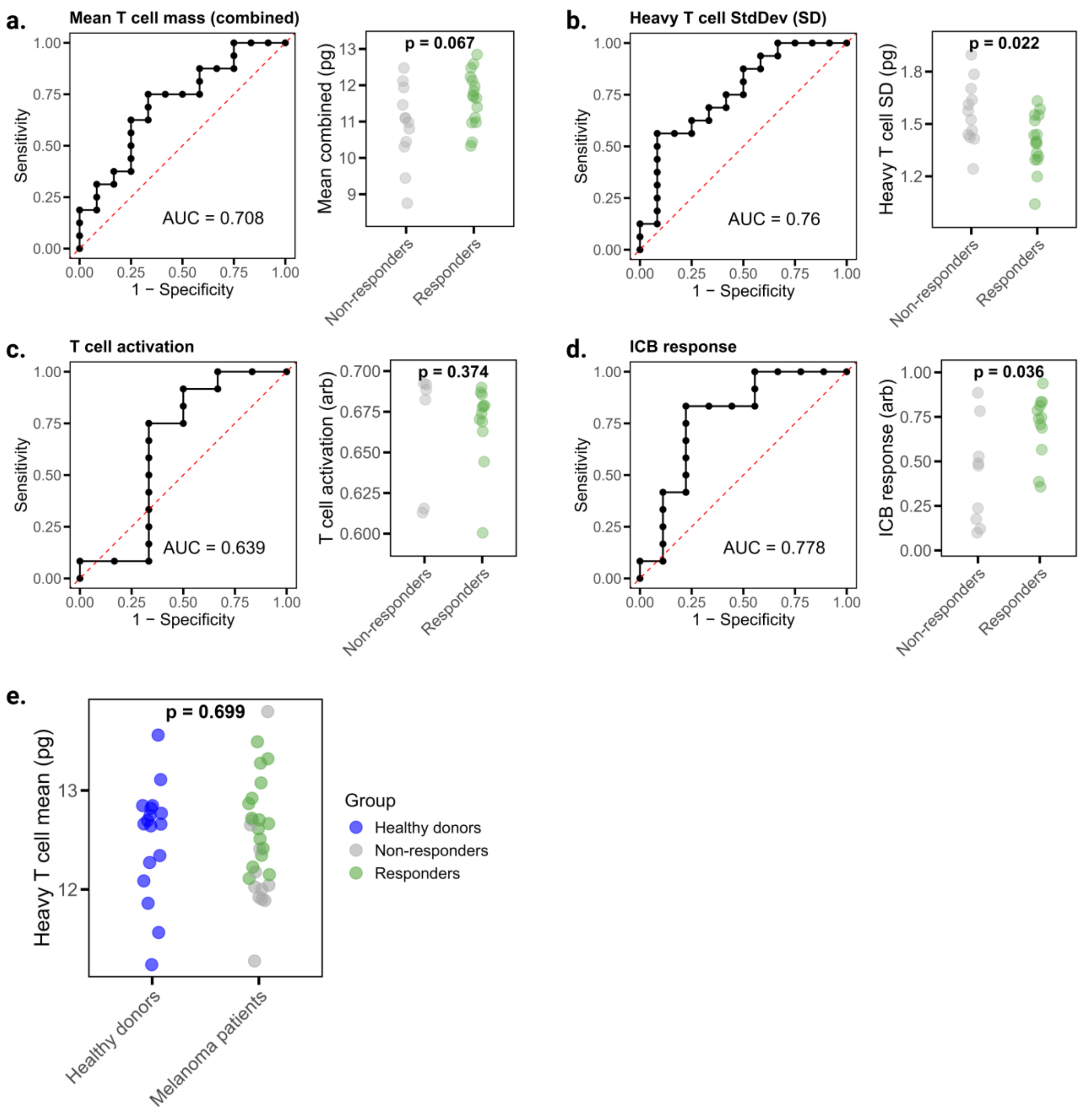


**Extended Data Fig.11 | Additional analysis of single-cell biophysical measurements and their association with ICB treatment response.**
**a.** Receiver operating characteristic (ROC) analysis (left) of the mean T cell mass across the full distribution (light and heavy populations) in predicting ICB response in the melanoma cohort (Fig. 3), with corresponding mean mass values compared between responders and non-responders (right). **b.** ROC analysis (left) of heavy T cell mass standard deviation (SD) in predicting ICB response, with SD values compared between responders and non-responders (right). **c.** ROC analysis (left) of ex vivo T cell activation in predicting ICB response, with corresponding activation values compared between groups (right). Activation was quantified by linear discriminant analysis (LDA) of two features: Earth Mover's Distance (EMD) metrics for single-cell mass and volume between unstimulated and anti-CD3/CD28–stimulated cells. **d.** ROC analysis (left) of T cell ICB response in predicting clinical ICB outcome, with corresponding LDA values compared between groups (right). ICB response was quantified by LDA of EMD metrics for mass and volume between cells stimulated with anti-CD3 alone versus anti-CD3 plus the ICB drug received by each patient. **e.** Comparison of mean heavy T cell masses in healthy donor PBMCs

versus melanoma patient samples. In all panels, p-values reflect Mann–Whitney U tests between responder and non-responder groups. AUC, area under the ROC curve.
